# Divide et Impera: Identification of Small-Molecule Inhibitors of HCMV Replication Interfering with Dimerization of DNA Polymerase Processivity Factor UL44

**DOI:** 10.1101/2020.02.06.938233

**Authors:** Hanieh Ghassabian, Federico Falchi, Veronica Di Antonio, Martina Timmoneri, Beatrice Mercorelli, Arianna Loregian, Giorgio Palù, Gualtiero Alvisi

## Abstract

Human cytomegalovirus (HCMV) is a leading cause of severe diseases in immunocompromised individuals, including AIDS and transplanted patients, and in congenitally infected newborns. Despite the availability of several antiviral drugs, their utility is limited by poor bioavailability, toxicity, and resistant strains emergence. Therefore, it is crucial to identify new targets of therapeutic intervention. The dimerization of HCMV DNA polymerase processivity factor UL44 plays an essential role in the viral life cycle being required for *ori*Lyt-dependent DNA replication. We validated the existence of UL44 homodimers both *in vitro* and in living cells by a variety of approaches, including GST pulldown, thermal shift, FRET and BRET assays. Dimerization occurred with an affinity comparable to that of the UL54/UL44 interaction, and was impaired by amino acid substitutions at the dimerization interface. Subsequently, we performed an *in-silico* screening to select 18 small molecules (SMs) potentially interfering with UL44 homodimerization. Antiviral assays using recombinant HCMV TB4-UL83-YFP in the presence of the 18 selected SMs led to the identification of four active SMs. The most active one also inhibited AD169 in plaque reduction assays, and impaired replication of an AD169-GFP reporter virus and its ganciclovir-resistant counterpart to a similar extent. As assessed by Western blotting experiments, treatment of infected cells specifically reduced viral gene expression starting from 48 h post infection, consistent with activity on viral DNA synthesis. Therefore, SMs inhibitors of UL44 dimerization could represent a new class of HCMV inhibitors, alternative to those targeting the DNA polymerase catalytic subunit or the viral terminase complex.

**IMPORTANCE:** HCMV is a ubiquitous infectious agent causing life-lasting infections in humans. HCMV primary infections and reactivation in non-immunocompetent individuals often result in life-threatening conditions. Antiviral therapy mainly targets the DNA polymerase catalytic subunit UL54 and is often limited by toxicity and selection of drug-resistant viral strains, making the identification of new targets of therapeutic intervention crucial for a successful management of HCMV infections. The significance of our work is in identifying the dimerization of the DNA polymerase processivity factor UL44 as an alternative antiviral target. We could show that full length UL44 dimerizes in a cellular context with high affinity and that such interaction could be targeted by small molecules, thus inhibiting the replication of several HCMV strains, including a drug-resistant mutant. Thus, our work could pave the way to the development of a new class of anti-HCMV compounds that act by targeting UL44 dimerization.

## INTRODUCTION

The β-*Herpesvirinae* member human cytomegalovirus (HCMV) is a major human pathogen, causing severe and life-threatening infections in immunocompromised patients (1) and in congenitally infected newborns (2). Herpesviruses are opportunistic double-stranded DNA viruses, whose genome transcription, replication, and packaging occur in the host cell nucleus (3). The molecular mechanisms involved in herpesvirus DNA replication and its regulation have been widely studied as they provide important models for the study of eukaryotic DNA replication and because viral enzymes involved in the process represent targets for antiviral therapy. HCMV DNA polymerase holoenzyme is a multi-functional enzyme that plays a key role during viral infection ensuring replication of the viral genome, and consists of the catalytic subunit UL54 and the processivity factor UL44 (4). Not surprisingly, the most widely antiviral agents used to fight HCMV infections target UL54 and are either nucleoside or pyrophosphate analogues such as ganciclovir (GCV) or foscarnet (PAA), respectively (5). However, long-term administration of these antiviral agents frequently leads to the selection of viral isolates with reduced drug susceptibility, due to mutations on either UL54 or on UL97, the viral kinase phosphorylating GCV (6, 7). Treatment with the recently approved Letermovir, which targets the viral terminase UL56 (8, 9), has been similarly shown to cause the selection of viral resistant strains (10, 11). Therefore, there is an urgent need to develop novel, potent anti-HCMV drugs which are directed against alternative targets, and possess activity against drug-resistant isolates.

UL44 is a 52-KDa protein essential for viral replication which binds to dsDNA and directly interacts with UL54, stimulating its activity by tethering the DNA polymerase holoenzyme to the DNA template (12–15). UL44 can be functionally and structurally divided in an N-terminal (residues 1-290) and a C-terminal domain (residues 291-433). The N-terminal domain has been successfully crystallized and retains all known UL44 biochemical properties (15, 16). However, the C-terminal domain, responsible for UL44 transactivation properties and its phosphorylation-dependent nuclear transport, is absolutely required for virus replication (17–21). UL44(1-290) adopts a C clamp-shaped structure and forms head-to-head dimers, wherein each monomer forms two topologically similar domains that share a central β-sheet and are connected by a long connector loop responsible for binding to UL54 (22, 23). The dimerization involves interaction of six main-chain-to-main-chain hydrogen bonds and extensive packaging of hydrophobic side chain at the interface and results in the formation of a central cavity, able to accommodate the viral DNA (24). Indeed, the UL44-dsDNA interaction depends on electrostatic interactions between the dsDNA backbone and basic residues located both on the central cavity and on a highly flexible gap loop not visible in the published crystal structure (24). Accordingly, the substitution of specific hydrophobic residues at the homodimerization interface of UL44(1-290) sufficient to impair dimerization, also reduces the DNA binding affinity of UL44 *in vitro* (16). Recent studies showed that dsDNA binding of UL44 is essential for HCMV DNA replication, as in the case of HSV-1 (25). Indeed, substitution of either basic gap loop residues, or of residues at the dimerization interface such as L86 and L87, which make extensive contacts with the hydrophobic residues along the dimer interface (16), caused a dramatic alteration of UL44 subcellular localization and DNA binding in cells, and completely abolished the ability of UL44 to trans-complement *ori*Lyt dependent DNA replication (23, 26, 27). Therefore, UL44 dimerization is important to stabilize the interaction between UL54 catalytic subunit and DNA and an alteration of this dimerization may block DNA synthesis and HCMV replication (28). In this context, UL44 dimerization thus emerges as a potential target for the development of novel antiviral approaches.

Protein-protein interactions (PPIs) are essential to all biological processes and can be modulated by small molecules (29–31), thus representing a large class of therapeutic targets and implying the possibility to impair viral replication and pathogenesis (32–34). Several inhibitors have reached the clinical trials thanks also to the development of computational and chemical technologies, alongside with experimental and virtual fragment screens used to define the druggability of PPIs (35, 36). Most of such inhibitors target PPIs in which partner proteins are characterized by short primary sequences at the interface (37, 38) and the hot-spots residues are concentrated in small binding pockets (39, 40). Recent studies have also reported the inhibition of protein dimerization, with a focus on proteins overexpressed in cancer or involved in viral replication (30, 41–43).

In this study, we aimed at identifying small molecules (SMs) that can hinder HCMV replication by interfering with UL44 homodimerization. To this end, we further characterized dimerization of full length UL44 both by assays *in vitro,* using GST pull-down and thermal shift assays (TSA), and in a cellular context by fluorescence and bioluminescence resonance energy transfer (FRET and BRET) assays. Subsequently, we used the UL44 homodimer structure as a template to screen about 1.3 million SMs from the ZINC database to identify 18 SMs potentially interfering with UL44 homodimerization via an *in-silico* screening. Our results clearly showed that one out of the 18 SMs tested (SM B3) could inhibit several HCMV strains, as assessed by plaque reduction, fluorescence reduction, and virus yield assays. As expected from a compound acting by inhibiting UL44 dimerization, SM B3 efficiently inhibited the replication of a HCMV GCV-resistant mutant and affected HCMV gene expression only starting from 48 h post infection (p.i.), causing a strong decrease of late gene pp28 expression at 72h and 96h p.i. Therefore, our results are consistent with a specific inhibition of B3 on viral DNA replication.

## MATERIALS AND METHODS

### Plasmid construction

Mammalian expression plasmids were generated using the Gateway^TM^ technology (Invitrogen). Entry clones pDNR-UL44(2-433), pDNR-UL44(2-433)L86A/L87A, pDNR-UL44(2-433)I135A, pDNR-UL44(2-433)Δloop, bearing the R165G/K167N/K168G triple substitution in the DNA–binding flexible gap loop, pDNR-UL44(2-433)ΔNLS, bearing the K428V/ K429A/K431L triple substitution in the NLS, pDNR-UL44(405-433), and pDNR-UL54(1220-1242) were previously described (20, 27, 44). Entry clones were used to generate C-terminal YFP fusion Mammalian expression vectors, following LR recombination reactions with the pDESTnYFP, pDESTnCFP, pDESTnRLuc Gateway compatible vectors (45), as described in (46). All constructs were confirmed by sequencing. Plasmids CFP-YFP, CFP and YFP were used as positive or negative controls for FRET experiments (47). Plasmids pDEST15-UL44 and pRSET-UL44 were used to express and purify from *E.coli* GST and His_6_-tagged proteins (4, 48).

### Proteins Expression and Purification

Recombinant GST-UL44 fusion protein and GST were purified from *Escherichia coli* BL21(DE3)/pLysS harbouring the respective plasmids pDEST15-UL44 and pDEST15, essentially as described in (49). Further details are available in the Appendix section.

### GST-pulldown assays

0.03 nmol of either GST or GST-UL44 were incubated with 0.12 nmol of either His_6_-UL44 or His_6_-UL44-L86/87A for 2 h at 4°C in binding buffer (25mM HEPES, pH 7.5, 12.5 mM MgCl_2_, 20% glycerol, 0.1% NP-40, 150 mM KCl, 0.15 mg/ml Bovine Serum Albumin [BSA], 1 mM DTT) under gentle shaking. Towards the end of incubation, 100 µl of Glutathione Sepharose 4 Fast Flow (GE Healthcare, #GE17-5132-01) were packed onto Poly-Prep chromatography columns (Biorad, #731-1550). The resin was equilibrated by addition of 1.5 ml of binding buffer. Samples were subsequently loaded onto the columns, columns washed with 6 ml of washing buffer (20 mM Tris-HCl, pH7.5, 100 mM NaCl, 0.1 mM EDTA, 0.5% NP-40). Bound complexes were eluted by addition of 400 µl of elution buffer (20 mM Tris-HCl, pH7.5, 100 mM NaCl, 0.1 mM EDTA, 0.5% NP-40 containing 19.5 mM Glutathione (Sigma Aldrich, #PHR1359)).

### Thermal Shift Assays (TSAs)

His_6_-UL44, His_6_-UL44-L86A/L87A (both at 5 μM final concentration), and Sypro orange (10 x final concentration) were diluted in Dilution Buffer (10 mM Na_2_HPO_4_, 10 mM NaH_2_PO_4_, 50 mM NaCl), and 20 μl of such mixtures were loaded on 96-well white plates (Microamp Optical 96-well Reaction Plate with Barcode, Applied Biosystems, #4306737), containing 5 μl of an appropriate 5x buffer, in duplicate. Buffers tested included three UL44-specific buffers: a DNA polymerase stimulation buffer (75 mM Tris-HCl, 150 mM KCl, 6.5 mM MgCl_2_, 1.67 mM β-mercaptoethanol, pH 8.0), an UL54-binding buffer (50 mM Hepes pH 7.5, 0.5 mM EDTA, 1 mM DTT, 150 mM NaCl, 4% Glycerol [v/v]), and a DNA-binding buffer (20 mM Tris-HCl, pH 7.5, 50 mM NaCl, 3 mM MgCl_2_, 0.1 mM EDTA, 0.1 mM DTT, 5% glycerol [v/v)]) (50–52). Additional buffers included either sodium citrate (pH 4.0, 5.0, 6.0, or 7.0), Tris-HCl (pH 7.0, 8.0 or 9.0), or bicine (pH 8.0 or 8.5) buffers either containing 50 or 250 mM NaCl (53). Plates were sealed with film (MicroAmp Optical Adhesive film, Applied Biosystems, #4311971) and centrifuged for 1’ at 700 rpm, before being heated from 22 to 95 °C, at 1°C/min using an ABI Prism 7000 Sequence Detection System (Applied Biosystems) set up as in (54), and data were acquired with a ROX emission filter. Data from TSA experiments were exported to Excel and analyzed to quantify the following parameters: Maximum Fluorescence, Maximum/Baseline fluorescence, dF/dT and T of Melting (Tm). Tm were calculated using the Boltzman Sigmoidal Fit Function with software GraphPad Prism (Graphpad Software Inc.) as described in (53).

### Cell culture

Human foreskin fibroblast (HFF), MRC5 and HEK 293T cells were maintained in Dulbecco’s modified Eagle’s medium (DMEM) supplemented with 10% (v/v) foetal bovine serum (FBS), 50 U/ml penicillin, 50 U/ml streptomycin, and 2 mML-glutamine and passaged when reached confluence. MCR5 cells were used up to passage number 30 and subsequently discarded.

### Fluorescence resonant energy transfer (FRET) acceptor photobleaching assays

The effects of specific substitutions on the ability of UL44 to self-interact or to bind the catalytic subunit UL54 were assessed by confocal laser scanning microscopy (CLSM) and fluorescence resonance energy transfer (FRET) acceptor bleaching as described in (47). For further details please refer to the Appendix section. HEK 293T cells were seeded in a 24-well plate onto glass coverslips (1×10^5^ cells/well) and the next day were transfected with 125 and 250 ng of YFP and CFP expression constructs, respectively, using Lipofectamine 2000 (Life Technologies) following the manufacturer’s recommendations. At 48 h post-transfection, cells were washed with PBS and fixed with 4% paraformaldehyde for 10’ at room temperature. After a wash with milliQ water, the samples were mounted on glass slides with Fluoromount G (Southern Biotech). Samples were imaged by CLSM using a Leica SP2 confocal laser scanning microscope (Leica) equipped with a 63× oil immersion objective. FRET efficiency was determined according to the following formula: FRETeff = [(EDpost − EDpre)/EDpost] × 100, where ED represents the emitted donor fluorescence before (EDpre) or after (EDpost) photobleaching of the acceptor fluorophore. Measurements of at least 20 cells from two independent transfections were used to calculate the box plots. Significance values were calculated by using the unpaired t test with the GraphPad Prism 6 software package (Graphpad Software Inc.). Images were processed using ImageJ (NIH).

### Bioluminescence resonance energy transfer (BRET) assays

BRET experiments were performed essentially as described previously using a reader compatible with BRET measurements (VICTOR X2 Multilabel Plate Reader, Perkin Elmer) (55). BRET saturation curves were generated using the GraphPad Prism software (Graphpad Software Inc.) by plotting each individual BRET ratio value to the YFPnet/RLuc signal, and interpolating such values using the one-site binding hyperbola function of GraphPad Prism (Graphpad Software Inc.) to calculate BRETmax (Bmax) and BRET_50_ (B_50_) values, indicative of maximum energy transfer and relative affinity of each BRET pair tested (56). For further details please refer to the Appendix section.

### Viruses, viral stocks preparation and titration

HCMV laboratory strain AD169 was obtained from ATCC (VR-53). Recombinant viruses AD169-GFP, expressing a humanized version of GFP under control of the HCMV IE promoter between open reading frames US9 and US10, as well as its GCV-resistant derivative AD169-GFP26, bearing the UL97 M460I substitution (57), were a generous gift by Manfred Marschall (Erlangen, Germany). Recombinant virus TB4-UL83-YFP, wherein YFP is fused to the C-terminus of the early-late tegument phosphoprotein pp65 (58) was kindly provided by Michael Winkler (Gottingen, Germany). All viral stocks were prepared and titred by immunological detection of IE1/2 proteins as described in (59) or by the plaque reduction method (see below). For details see the appendix section.

### Analysis of UL44 dimerization interface

The crystallographic structure of UL44 homodimer (16) was downloaded from the Protein Data Bank(pdb code: 1T6L). Only the A chain was extracted from the complex. The structure was optimized with the Protein Preparation Wizard tool of the Schrodinger suite (Schrödinger). The research of potentially druggable sites at the dimerization interface of UL44(1-290) was assessed with software Sitemap (60), using default settings apart from *grid resolution* which was set as *fine*.

### Virtual screening database preparation

A 3D molecular database was built with the Schrodinger suite (Schrödinger) starting from 2D structures taken from the ZINC database (www.zinc.docking.org). A total number of about 3.6 million compounds were selected (Vendors: Asinex, Chembridge, Princeton, NCI and ZINC natural) and downloaded for this study. 2D structures were converted into 3D structures and stereoisomers were generated with the Ligprep function of the Schrodinger suite (Schrödinger). Moreover, for each entry all the possible ionization states at pH 7.0 ± 2.0 and tautomers were generated with Epik (Schrödinger). The obtained database consisted of about 5 million compounds. In order to retrieve the most drug-like compounds, ADMET properties of each molecule in the database were predicted with Qikprop (Schrödinger) and compounds were filtered as described in (61) using a “soft Lipinsky rule” (Molecular weight ≤ 600, Rotable bonds ≤ 10, Number of H-bond acceptors ≤ 10, Number of H-bond donors ≤ 5, Number of chiral centers ≤ 2, QplogPo/w ≤ 6). Finally, the number of compounds was reduced to about 1.3 million by using the PPI-HitProfiler software with the “soft” mode (CDithem; http://www.cdithem.fr).

### Virtual screening

For the docking stage, a receptor grid was built on the UL44 structure prepared as above. The grid was centered on the position of residue M116 and receptor grid generation default settings were applied. By using this grid, the database was docked with the High throughput virtual screening (HTVS) scoring function of the Glide software (Schrödinger). After this run, we selected 50,000 compounds on the basis of the HTVS docking score. Selected molecules were submitted to a second run of docking using the standard precision (SP) scoring function and only the 5,000 top-ranked compounds were selected. Finally, selected molecules were submitted to a run of docking using the extra precision (XP) scoring function and only the 500 top-ranked compounds were selected. After docking, we further narrowed down the number of selected compounds to 18 on the basis of commercial availability, visual inspection, and by a cluster analysis performed with Tanimoto on the basis of the Molprint2D fingerprints of each molecule (62).

### Preparation of small molecules (SMs) and Ganciclovir (GCV) stocks

Ganciclovir (GCV; Selleckchem, S1878) and small molecules (SMs), with a > 90% purity assessed by liquid chromatography mass spectrometry and high-performance liquid chromatography (Vitas-M Laboratory), were resuspended in 100% DMSO to obtain 20 mM and 50 mM stocks, respectively, and stored at -20°C protected from light.

### Antiviral compounds testing

For identification of SMs active on HCMV replication, MRC5 cells (1.5×10^4^ cells/well) were seeded in 96-well special optics black microplates (Corning, CLS3614) and incubated overnight at 37°C, 5% CO_2_ and 95% humidity. The next day, cells were infected for 1 h at 37 °C in 100 μl/well of DMEM containing TB4-UL83-EYFP at an MOI of 0.03 infectious units/cell. Cells were subsequently washed, and media containing either 0.5% DMSO or two different concentrations (100 and 10 μM) of SMs with a 0.5% DMSO final concentration was added. 50 μM GCV was included as a positive control for inhibition of viral replication. Mock infected cells served as a reference for calculation of background fluorescence. After addition of SMs, the plates were further incubated at 37°C, 5% CO_2_ and 95% humidity. Every day, cell confluence and morphology, as well as CPE and the presence of precipitates were evaluated by light microscopy. Fluorescence signals were visualized on an inverted fluorescent microscope (Leica, DFC420 C). Seven days p.i., cells were washed once with PBS and lysed with luciferase lysis buffer (25 mM glycylglycine, 15 mM MgSO_4_, 4 mM EGTA, 0.1 % Triton X-100, pH 7.8). Plates were immediately frozen at -20°C, thawed at RT, and fluorescent signals relative to each condition were acquired using a reader compatible with fluorescence measurements (VICTOR X2 Multilabel Plate Reader, Perkin Elmer) equipped with a fluorimetric excitation filter (band pass 485 ± 14 nm) and a fluorimetric emission filter (band pass 535 ± 25 nm). After background subtraction, data were normalized to solvent-treated controls and analyzed with Graphpad Prism (Graphpad Software Inc.). The screening was performed three times, and each plate included at least two wells treated with the same compound, as well as at least 12 wells treated with solvent only.

### Fluorescent reduction assays (FRA)

To calculate the effective dose 50 (ED_50_) of each compound against TB4-UL83-EYFP virus by means of FRA, MRC5 cells were seeded, infected, treated, and processed as above, using increasing concentrations of each SMs and GCV (range between 0.02 and 100 μM). Mock infected cells served as a reference for calculation of background fluorescence. After background subtraction, data were normalized to solvent-treated controls and analyzed with Graphpad Prism (Graphpad Software Inc.). The experiments were performed six times, and each plate comprised at least two wells treated with the same compound, as well as at least 14 wells treated with solvent only. To calculate the ED_50_ of each compound against AD169-GFP virus and its GCV-resistant AD169-GFP26 counterpart by the means of FRA, MRC5 cells were seeded in 12-well plates (1.8×10^5^ cells/well) in 1 ml DMEM supplemented with 10% (v/v) FBS, 50 U/ml penicillin, 50 U/ml streptomycin and 2 mM L-glutamine, and incubated overnight at 37°C, 5% CO_2_ and 95% humidity. The next day, cells were infected for 2 h at 37 °C in 1 ml/well of DMEM at MOI of 0.05 infectious units/cell as described in (57). Subsequently, cells were washed with 2 ml of PBS, and 1 ml of medium containing either 0.5% DMSO or increasing concentrations (from 0.001 to 100 μM) of each compound with a 0.5% DMSO final concentration was added. Every day, cell confluence and morphology, as well as CPE and the presence of precipitates were evaluated by light microscopy, whereas fluorescence signals were visualized on an inverted fluorescent microscope (Leica, DFC420 C). Seven days p.i., supernatants were collected, cleared from cells and debris by centrifugation for 5’ at 700 rpm, and stored at -80°C until used for virus yield assays (VIAs) as described below. Cells were washed with 2 ml of ice cold PBS and lysed in 200 μl of GFP lysis buffer (25 mM Tris-HCl, pH 7.8, 2 mM DTT, 2 mM trans-1,2-diaminocyclohexane-N,N,N,N-tetraacetic acid, 1% Triton X-100, 10% glycerol [v/v]). Plates were further incubated 10’ at 37°C in a humidified incubator, before being incubated for 30’ at RT with shaking at 225 rpm. Samples were centrifuged for 5’ at 4°C at 13,000 rpm and 100 μl of cleared lysates were transferred to black bottomed 96-well plates (Costar, #3916). Fluorescent signals were acquired and analyzed as described above.

### Virus yield assays (VIAs)

To determine the ED_50_ of each compound against AD169-GFP virus and its GCV-resistant AD169-GFP26 counterpart by the means of VIAs, MRC5 cells were seeded in clear flat bottom 96-well tissue culture plates (1.5×10^4^ cells/well) with low evaporation lids (Falcon, #353072). The next day, medium was replaced with serial dilutions of supernatants containing AD169-GFP or AD169-GFP26 virus grown in the presence of inhibitory compounds. One week later, virus yield relative to each condition was calculated and expressed as 50% Tissue Culture Infectious Dose (TCID_50_)/ml using the Spearman and Karber algorithm as described in (63).

### Plaque reduction assays

For plaque reduction assays (PRA) human foreskin fibroblast (HFF) cells were seeded in 24-well plates (2×10^5^ cells/well). The following day cells were infected at 37°C with 70 Plaque Forming Unit (PFU) of HCMV AD169 per well in DMEM containing FBS 5%. At 2 h p.i., the inocula were removed, cells were washed, and media containing various concentrations of each compound, 5% FBS, and 0.6% methylcellulose were added. All compound concentrations were tested at least in triplicate. After incubation at 37°C for 10-11 days, cell monolayers were fixed, stained with crystal violet, and viral plaques were counted.

### Cell cytotoxicity assays

MRC5 cells (1.5×10^4^ cells/well) were seeded in clear flat bottom 96-well tissue culture plates with low evaporation lids (Falcon, #353072) in duplicate. After 24 h, cells were treated with different concentrations of GCV or SMs, or solvent only. A number of wells containing only DMEM and no cells were also included for background correction. Seven days post-treatment, cells were processed for measurement of cell metabolic activity using 3-(4,5-Dimethyl-2-thiazolyl)-2,5-diphenyl-2H-tetrazoliumbromid (MTT; Applichem, #A2231) following the manufacturer’s recommendations. See Appendix for details.

### Analysis of HCMV gene expression

MRC5 cells were seeded on 6-well flat bottom plates (6×10^5^/well) with low evaporation lid (Falcon, #353046). The following day, cells were either mock infected or infected with HCMV (strain AD169) at MOI of 2 infectious units/cell in DMEM at 37°C. One hour p.i., cells were washed twice with PBS and medium containing either GCV (16 μM), B3 (50 μM) or solvent only (0.5% DMSO) was added to each well. At different times p.i., cells were washed twice with PBS and lysed on ice with 250 μl of RIPA buffer containing protease inhibitors (Tris-HCl 50 mM, pH 7.4, 150 mM NaCl, 1% Triton X-100 (v/v), 1% sodium deoxycholate, 0.1% SDS, 1 mM EDTA, 17.4 μg/ml phenylmethylsulfonyl fluoride, 2 μg/ml aprotinin, and 4 μg/ml leupeptin). The protein content in each sample was quantified using the Micro BCA Protein Kit assay (Thermo Scientific).

### Western-blot assays

Fractions derived from GST-pulldown assays or 30 μg of infected cell lysates were analyzed by SDS page/Western blotting as described previously (49). The following antibodies, diluted in PBS containing 0.2% Tween20 and 5% milk (w/v), were used: *α*-His_6_ mAb (Sigma Aldrich, #H-1029; 1:2,500), *α*-IE1&2 mAb (Virusys Corporation, #CA003-1; 1:10,000), *α*-UL44 mAb (Virusys Corporation, #P1202-1; 1:100); *α*-pp65 mAb (Virusys Corporation, #P1251; 1:2,000); *α*-pp28 (Abcam, #ab6502; 1:10,000), rabbit *α*-GADPH pAb (Santa Cruz Biotech, #sc-25778; 1:5,000); mouse *α*-β-Actin mAb (Sigma Aldrich, # A5316 1:5,000); goat *α*-mouse (Santa Cruz Biotech, #sc-2055; 1:10,000) and *α*-rabbit (Sigma Aldrich, #A6154; 1:10,000) immunoglobulin Abs conjugated to horseradish peroxidase. Signals were acquired using an imaging system (Alliance Mini, Uvitech) and quantified using Image J (NIH).

## RESULTS

### Full length UL44 dimerizes *in vitro* and a few residues are essential for self-interaction

All data relative to UL44 dimerization *in vitro* rely on the deletion mutant UL44(1-290), which lacks the highly unstructured C-terminal domain of the protein. Therefore, we investigated whether full length UL44 similarly dimerizes *in vitro* by means of GST-pulldown assays (Figure 1A). Our results indicate that His_6_-UL44 could be specifically pulled down by GST-UL44 but not by GST alone (Figure 1A *left* and *middle panels*). On the other hand, the fraction of His_6_-UL44-L86A/L87A pulled down was clearly reduced, indicating impaired ability to associate with GST-UL44, and thus further supporting the validity of the previously published crystal structure (Figure 2A, *right panel*). Since single amino acids substitutions impairing self-association of multimeric proteins are known to decrease protein stability (41), we decided to further corroborate our findings by comparing the stability of His_6_-UL44 and its L86A/L87A derivative by thermal shift assays (TSAs). To this end, we initially performed a small-scale screening to identify the most suitable conditions to assess UL44 stability by TSA (Supplementary Figure S1). Our data indicated that His_6_-UL44 is significantly less stable as compared to lysozyme under the 22 conditions tested (Tm=48.0±3.5°C and 73.2±1.8°C, respectively; see Supplementary Figure S1AD), and its stability is strongly dependent on pH (Supplementary Figure S1EF), but not on salt concentration (Supplementary Figure S1GH). Such analysis allowed us to identify the 3 most suitable buffer conditions to assess UL44 stability by TSA (Supplementary Figure S1IJ). We next compared the effect of the L86A/L87A double substitution on UL44 stability. Our results showed that His_6_-UL44 is significantly more stable than His_6_-UL44-L86A/L87A (Tm=52±0.3°C *vs*. 42.4±1.4°C, respectively, in the three buffers tested, see Figure 2B), consistent with the involvement of L86 and L87 in protein dimerization.

**Figure 1.**
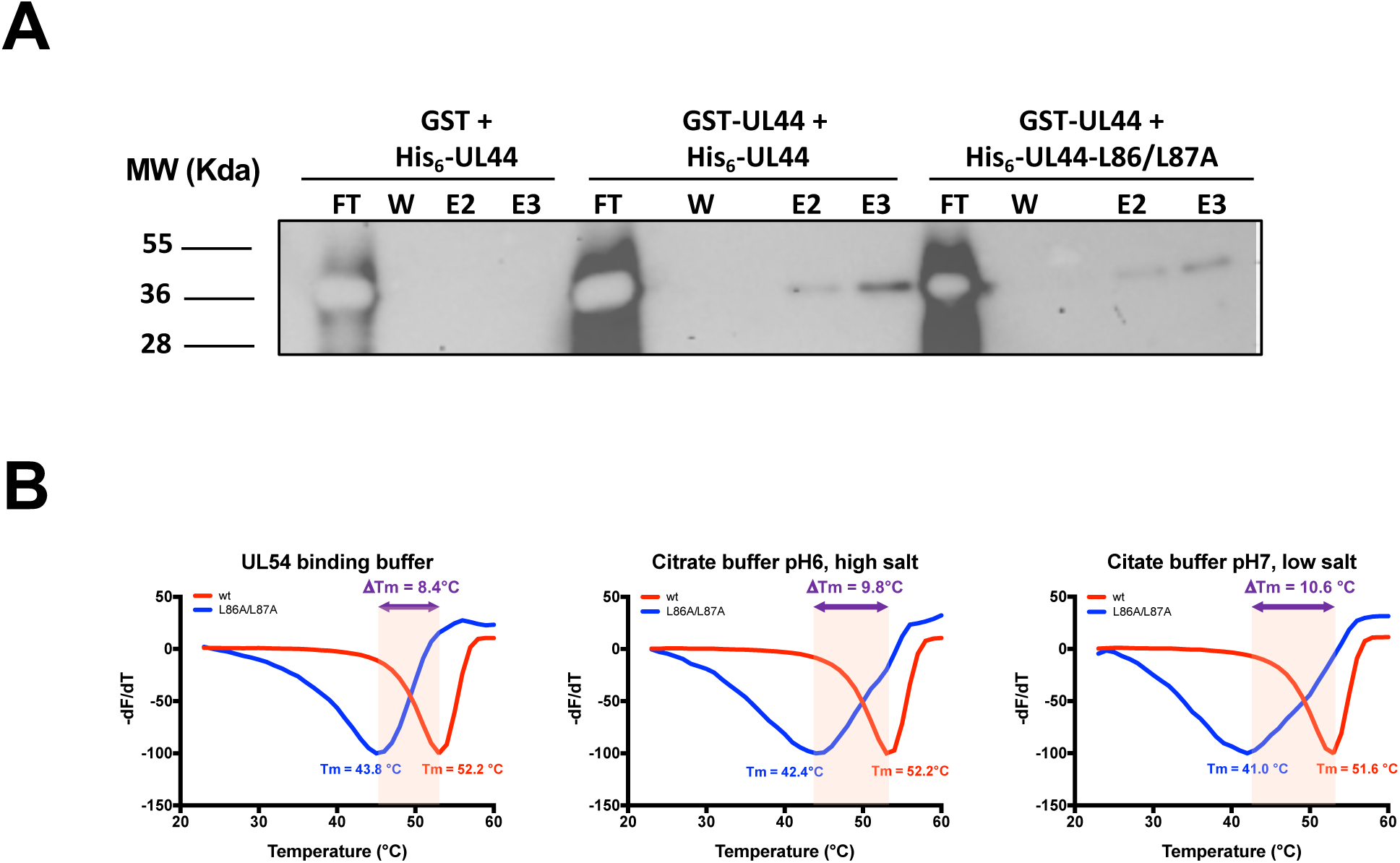
Characterization of full length UL44 self-interaction *in vitro*. *(A)* The indicated GST and His_6_ fusion proteins were premixed before being subjected to GST pulldown and SDS/PAGE-Western blotting using specific antibodies directed against the His_6_-tag. FT, flowthrough; W, wash, E, elution. *(B)* Melting curves obtained for different UL44 variants in thermal shift assays (TSA). The indicated His_6_-UL44 fusions were subjected to TSA experiments in the specified buffers as described in the Materials and Methods section. Data plotted are the –dF/dT against the Temperature of experiments performed in duplicate.

**Figure 2.**
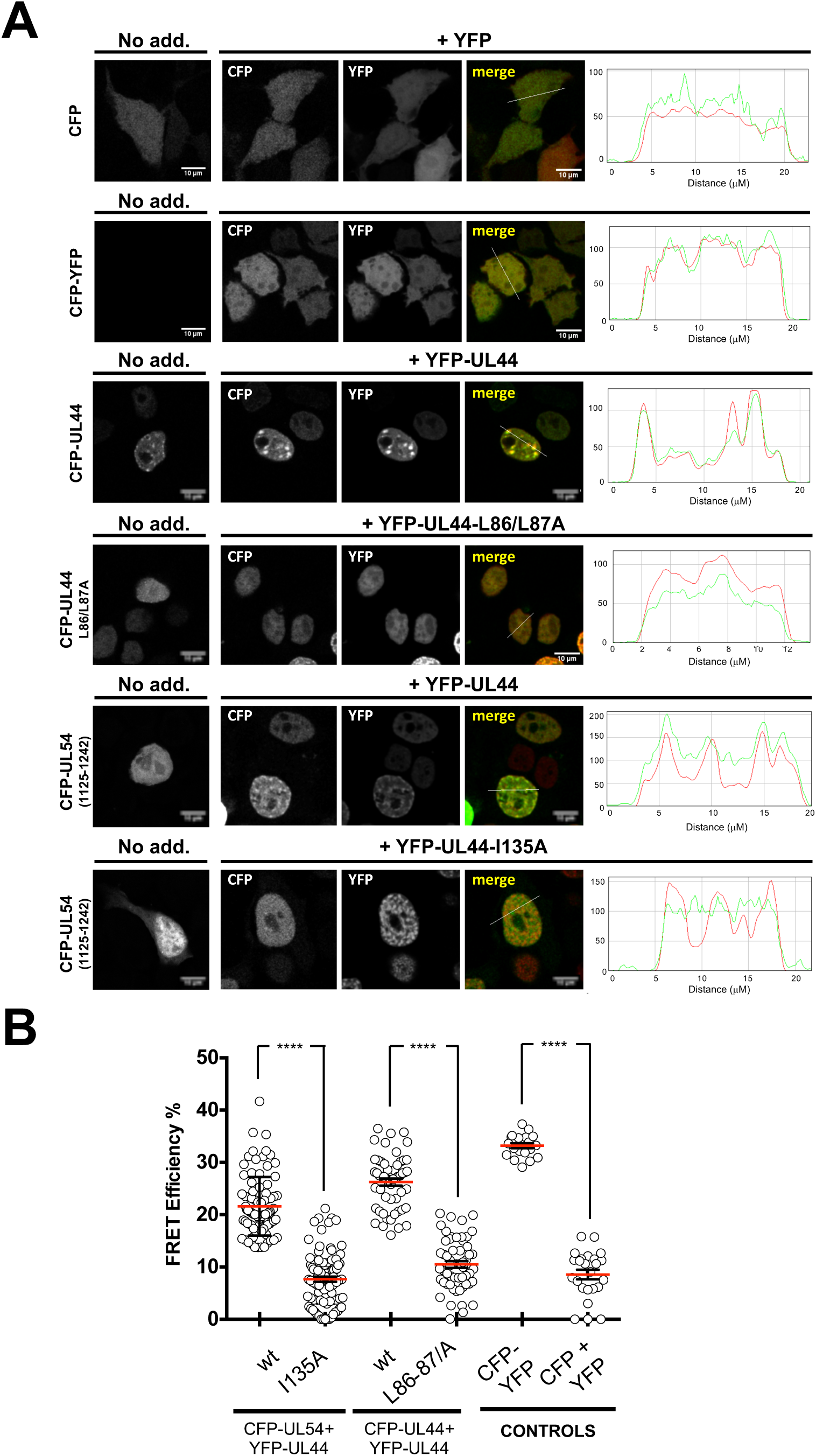
Characterization of UL44 self-interaction in cells. HEK 293T cells were transfected with the indicated expression plasmids and at 48 h post-transfection, cells were fixed and processed for IF, before being analyzed by CLSM to determine the subcellular localization of the indicated fusion proteins. *(A)* The subcellular localization of CFP fusions as expressed alone (*left panels*) or in the presence of YFP fusions (*middle panels*) is shown, together with the relative RGB profiles (*right panels*). *(B)* Cells micrographs such as those shown in *(A)*, were subjected to FRET acceptor photobleaching to calculate FRET efficiency, as described in the Materials and Methods section. Data shown are the mean and standard deviation relative to at least 20 cells from two independent experiments, along with the p-value relative to the Student’s t-test relative to the indicated groups of FRET pairs; ****: p ≤ 0.0001.

### UL44 dimerizes *in cells* and a few residues are essential for self-interaction

We have previously shown that different UL44 variants can reciprocally influence their subcellular localization by CLSM analysis of GFP and DsRed2 fusions (23, 26), suggesting that the protein can also exist as a dimer in a cellular context. However, interpretation of such relocalization assays could be complicated by experimental artefacts due to DsRed2 oligomerization (64). We therefore decided to further corroborate such findings by fluorescent energy transfer (FRET) assays. To this end, HEK 293T cells were transfected with plasmids mediating the expression of CFP-UL44 or its L86A/L87A and I135A derivatives, expected to be impaired in dimerization and in binding to UL54, respectively. Cells were also co-transfected with plasmids expressing YFP-tagged UL44 and the C-terminus of UL54 bearing the UL44-binding domain (amino acids 1125-1242). Cells were also transfected to express CFP and YFP individually, as negative controls, or a CFP-YFP fusion protein, as a positive control. We initially assessed the subcellular localization of each fusion protein when expressed individually (Figure 2A, *left panels*). As expected, both CFP and CFP-YFP localized with a diffused pattern throughout the cell. On the other hand, CFP-UL44 localized in the cell nucleus, with a punctate pattern, dependent on its ability to bind to dsDNA (27). The dimerization defective CFP-UL44-L86/L87A still localized within the cell nucleus, due to the presence of a functional NLS, but with a diffuse pattern, consistent with its reduced DNA-binding ability (23). CFP-UL54(1125-1242), similarly containing a functional NLS, but devoid of DNA-binding properties, localized to the cell nucleus with a diffuse pattern (44). Coexpression of CFP- and YFP-tagged UL44 resulted in the two proteins colocalizing into the cell nucleus with evident accumulation in punctate structures, while co-expression of CFP- and YFP-tagged UL44-L86A/L87A derivatives led to co-localization within the nucleus, but with a diffuse pattern (Figure 2A, *middle panels*). Importantly, upon co-expression with YFP-UL44, CFP-UL54(1125-1242) relocalized to nuclear punctate structures, while its localization mainly remained diffused upon co-expression with YFP-UL44-I135A. Our FRET analysis (Figure 2B), revealed a high FRET efficiency for the CFP-YFP fusion protein (33.2±0.5; n=20), but not for the co-expression of CFP and YFP as individual proteins (8.6±0.9; n=24). On the other hand, co-expression of CFP- and YFP-tagged UL44 resulted in a significantly higher FRET efficiency (23.3±0.7; n=54) as compared to their L86A/L87A counterparts (10.5±0.6; n=57). Similarly, expression of CFP-tagged UL54 in the presence of UL44 resulted in a significantly higher FRET efficiency (21.6±0.6; n=89) as compared to the FRET efficiency in the presence of an UL44 mutant bearing the I35A substitution, which prevents the interaction with UL54 *in vitro* (7.7±0.5; n=97).

Clearly, these data indicate that full length UL44 exists as a dimer in a cellular context, and dimerization relies on residues identified in the crystal structure of its N-terminal domain.

### UL44 self-interacts in cells with an affinity comparable to that of the UL44-UL54 interaction

Previous studies proposed the UL44-UL54 interaction as a potential target for therapeutic intervention and identified SMs inhibiting HCMV replication by interfering with holoenzyme formation (48, 65). Our data suggest that UL44 dimerization surface might be a similarly suitable target for the development of antivirals. *In vitro*, the affinity of UL44 self-interaction has been reported to be similar to that of the UL44-UL54 interaction, although using different methods and thus not being directly comparable (16, 22). We therefore decided to compare the affinity of such interactions using a bioluminescent resonant energy transfer (BRET) assay that we recently developed to study respiratory syncytial virus (RSV) matrix protein self-association in living cells (55). HEK 293T cells were transfected to express either RLuc-UL44 or RLuc-UL44(405-433) either in the absence or in the presence of YFP-UL44, and 48 h later cells were processed for BRET analysis. The RLuc-UL44(405-433) fusion protein contains an NLS, responsible for nuclear targeting of the protein, but lacks the UL44 N-terminal domain responsible for all known biochemical activities, including the ability to self-associate, and therefore represents an ideal negative control for BRET assays. Importantly, under all conditions tested, both RLuc (Supplementary Figure S2A) and YFP fusions (Supplementary Figure S2B) were expressed to comparable levels. When individually expressed, RLuc-UL44 generated a BRET value of 0.31±0.01, similar to the 0.32±0.01 BRET value calculated for Rluc-UL44(405-433) and also corresponding to the background signal. Importantly, co-expression of YFP-UL44 increased the RLuc-UL44 BRET value to 0.67±0.05, resulting in a BRET ratio of 0.36±0.10. On the other hand, BRET value of RLuc-UL44(405-433) was not increased by co-expression with YFP-UL44 (0.35±0.01), resulting in a BRET ratio of 0.03±0.00 (Supplementary Figure S2CD). These data suggest that BRET can be used to quantify UL44 self-interaction in living cells. Importantly, when BRET saturation experiments were performed by transfecting cells with a fixed amount of BRET donor plasmid RLuc-UL44 or RLuc-UL44(405-433) in the presence of increasing amounts of BRET acceptor plasmid YFP-UL44, the BRET ratio relative to RLuc-UL44 and YFP-UL44 promptly reached saturation, whereas the BRET ratio relative to RLuc-UL44(405-433) and YFP-UL44 did not (Figure 3A). Data fitting allowed us to calculate the Bmax value, corresponding to the maximal BRET ratio obtainable for a BRET pair, and the B_50_ value, corresponding to the ratio between BRET acceptor and donor sufficient to generate a BRET ratio corresponding to half of the Bmax, and indicative of the affinity of UL44 self-interaction in living cells. Similar BRET saturation experiments were performed for RLuc-UL54/YFP-UL44 in order to compare UL44 dimerization and UL44 binding to UL54 (Figure 3B). Importantly, UL44 appeared to self-interact with an affinity similar to that of the UL54-UL44 interaction, as suggested by the very similar Bmax (0.64±0.09; n=8 and 0.52±19.56; n=5, respectively) and – most importantly –B_50_ values (19.04±13.69; n=8 and 19.96±13.88; n=5) values (Figure 3C-F and Table I). The same approach was then used to study the effect of specific UL44 substitutions on its homodimerization and on UL54 binding (see Table I). Importantly, substitutions at the UL44 dimerization interface (F121A, L86A/L87A) specifically increased the B_50_ value relative to UL44 dimerization but not that relative to binding to UL54, implying that the corresponding proteins were not misfolded (see Figure 3D, E and Table I). On the other hand, the I135A substitution specifically increased the B_50_ value relative to binding to UL54 but not that relative to UL44 dimerization. As expected, the Δloop and ΔNLS substitutions did not affect either process. Thus, UL44 self-interacts in living cells with an affinity comparable to that of the UL44-UL54 interaction and single amino acid substitutions at the dimerization interface strongly impair the process.

**Figure 3.**
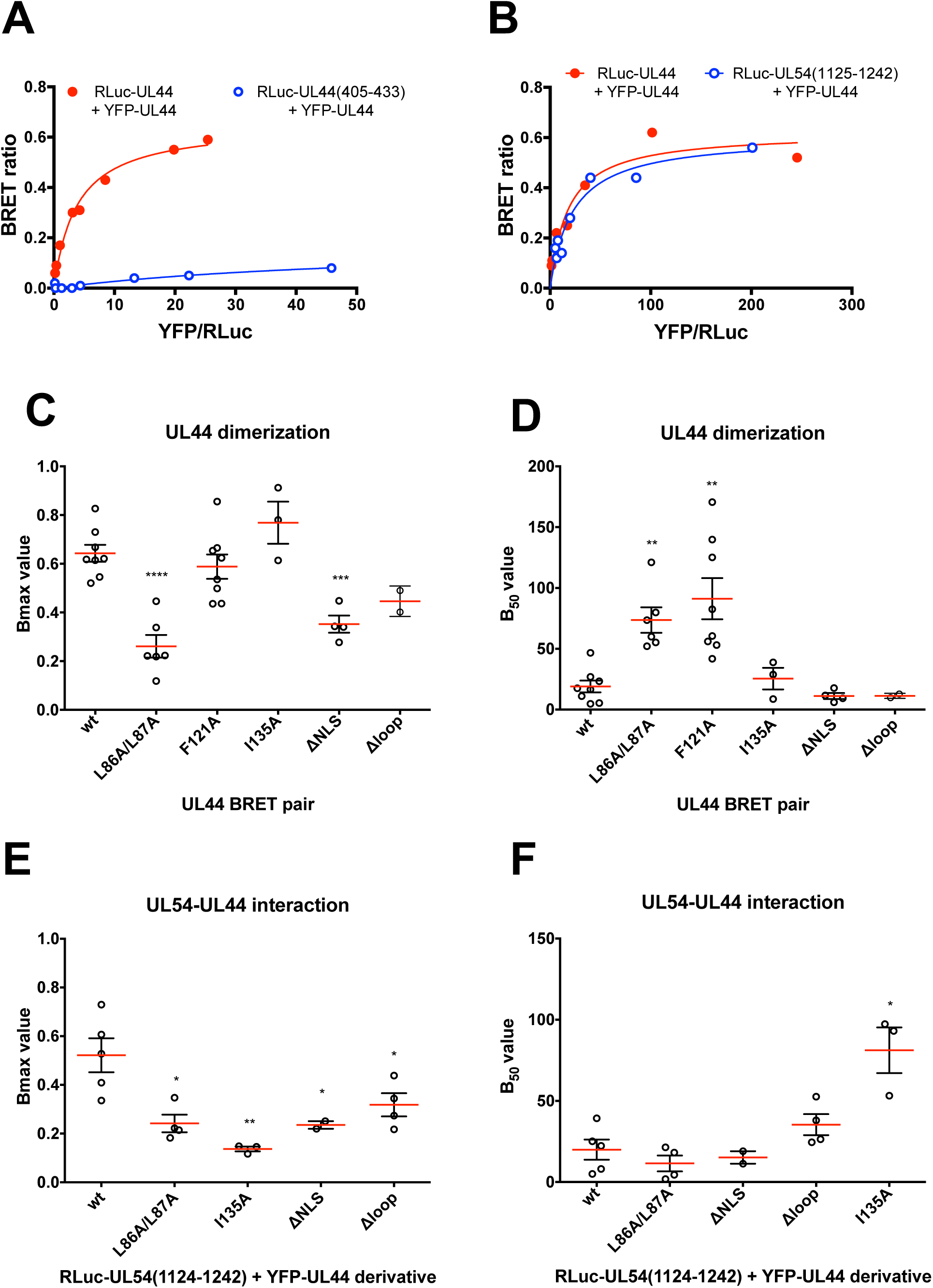
The affinity of UL44 to itself in living cells is similar to that of the interaction of UL44 to UL54. (A, B) HEK 293T cells were transfected to transiently express the indicated RLuc-UL44 fusion proteins either in the absence or in the presence of increasing amounts of plasmid YFP-UL44 and processed for BRET measurements as described in the Materials and Methods section. The BRET ratio relative to the indicated BRET pair was plotted against the YFPNet/RLuc ratio and data used to calculate the Bmax and B_50_ values. Representative data from at least three independent experiments are shown. (C-F) HEK 293T cells were transfected to transiently express either the indicated RLuc-UL44 (C, D) or RLuc-UL54 (E, F) fusion proteins, in the presence of increasing amounts of plasmids expressing the indicated YFP-UL44 derivatives, to generate BRET saturation curves such as those shown in (B) and calculate the Bmax (C, E) as well as the B_50_ (D, F) values. Data shown are single measurements, mean and standard deviations of the mean of at least three independent experiments, along with the p-value relative to the Student’s t-test the indicated groups of BRET pairs; *: p ≤ 0.05, **: p ≤ 0.005

**Table I.** Summary of BRET saturation experiments.

| UL44 derivative | RLuc-UL44 + YFP-UL44 |  | RLuc-UL54 + YFP-UL44 |  |
| --- | --- | --- | --- | --- |
|  | <u>B<sub>max</sub></u> | <u>B<sub>50</sub></u> | <u>B<sub>max</sub></u> | <u>B<sub>50</sub></u> |
| Wild-type | 0.64±0.09 (8) | 19.0±13.7 (8) | 0.52±19.56 (5) | 20.0±13.9 (8) |
| L86A/L87A<sup>1,5</sup> | 0.26±0.11 (6) | 73.7±25.7 (6) | 0.24±0.07 (4) | 11.5±9.8 (4) |
| I135A<sup>2</sup> | 0.77±0.15 (3) | 25.5±15.4 (3) | 0.14±0.02 (3) | 81.2±24.3 (3) |
| F121A<sup>1</sup> | 0.59±0.14 (8) | 91.2±47.8 (8) | N.A. | N.A. |
| ΔNLS<sup>3,5</sup> | 0.35±0.07 (4) | 11.2±4.9 (4) | 0.24±0.02 (2) | 15.1±5.4 (2) |
| Δloop<sup>4,5</sup> | 0.45±0.06 (2) | 11.3±2.1 (2) | 0.32±0.10 (4) | 34.4±13.0 (4) |

### An *in-silico* screening identifies small molecules potentially interfering with UL44 dimerization

We aimed at identifying small molecules (SMs) interfering with UL44 self-interaction. To this end, the published crystal structure of UL44 (Figure 4A) was analyzed with SiteMap to identify top-ranked potential receptor binding sites on the protein monomer. Our analysis identified 3 sites as potential receptor binding sites, one of which sites is located at the interface between the two monomers (Figure 4B). Such pocket has a druggability score (Dscore) of ≅ 0.8 according to SiteMap (0.797) and it is therefore potentially druggable (60). The key residues seem to be L87 and especially L86, which are located in a cavity formed by the residues F121, M123, M116, L93, C117, K101, T100, A118, L99, S96, D98 and P119 of the other monomer. Therefore, the center of the grid built for the docking calculation was set on M116 and such structure was used to screen about 1.3 million compounds from the ZINC database using the Glide software (Figure 4C). Eighteen of these molecules were purchased and used for further assays (Supplementary Figure S3 and Supplementary Tables I and II).

**Figure 4.**
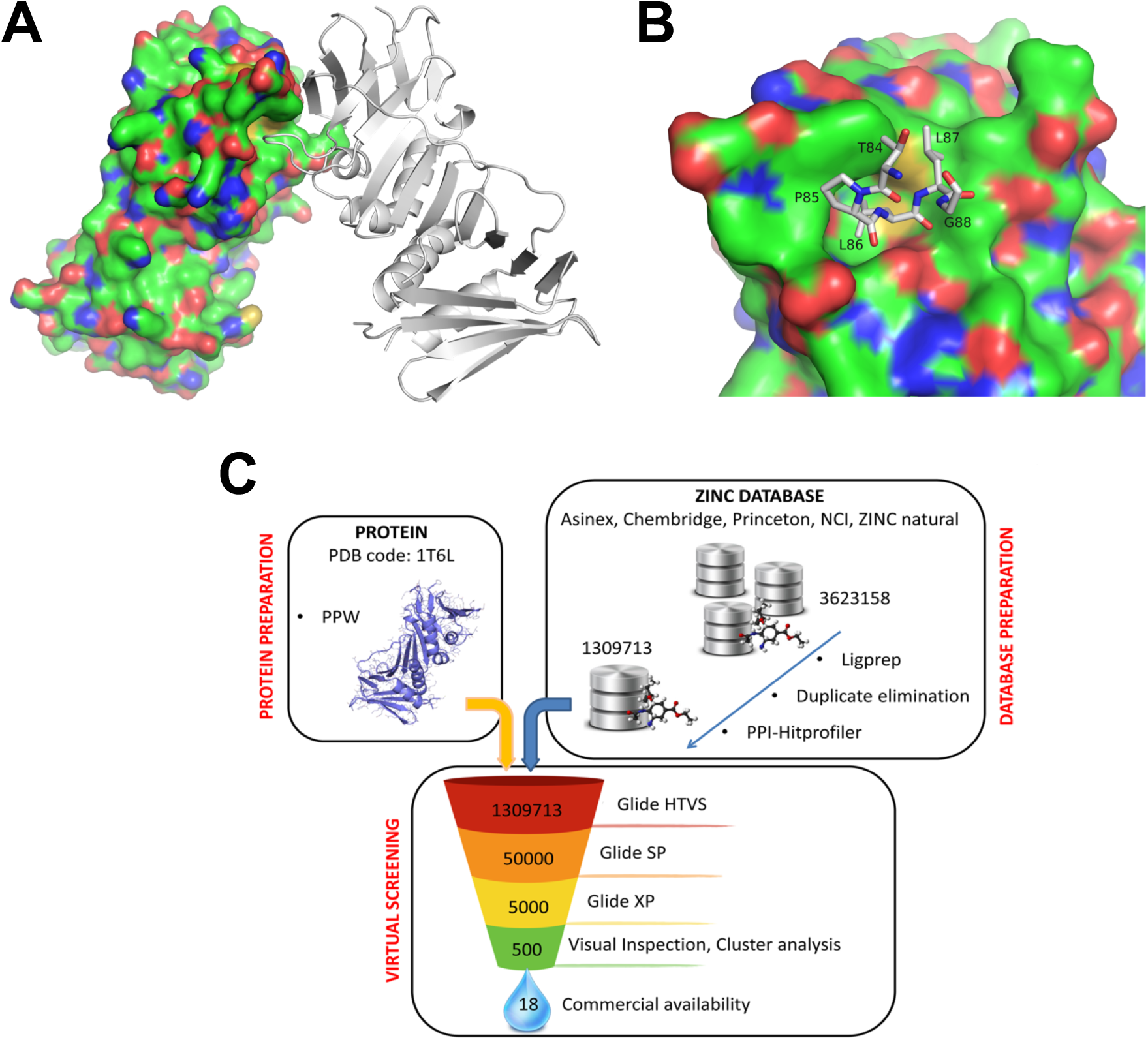
An *in-silico* screening to identify SMs inhibiting UL44 dimerization. (A) Graphic representation of UL44(1-290) homodimers. One monomer is represented as surface, the other one as ribbons, with residues involved in the dimerization being shown as sticks. (B) One monomer is represented as surface, with residues involved in the dimerization from the other monomer being shown as sticks. (C) A virtual screening leads to identification of SMs potentially disrupting UL44 homodimerization. The Glide software was used to dock molecules to the interface of the two monomers (PDB code: 1T6L). Three rounds of screening by were performed using the High throughput virtual screening (HTVS), Standard Precision (SP) and Extra Precision (XP) docking settings. After each docking, round the top-ranked molecules in term of docking score were selected for the following round. The resulting 500 molecules were further reduced by visual inspection, cluster analysis and, on the basis of their commercial availability, 18 compounds were purchased and tested for their ability to disrupt UL44 dimerization.

**Table II.** Summary of FRA and VYA for AD169-GFP and AD169-GFP26.

|  | FRA (*n*=5) ED<sub>50</sub> (μM)<sup>1</sup> |  |  | VIA (*n*=3) ED<sub>50</sub> (μM) |  |  |
| --- | --- | --- | --- | --- | --- | --- |
|  | AD169-GFP | AD169-GFP26 | ratio | AD169-GFP | AD169-GFP26 | Ratio<sup>2</sup> |
| GCV | 2.6±0.8 | 20.0±4.3 | 9.7±2.3 | 0.5±0.5 | 4.5±4.5 | 34.7±30.2 |
| B3 | 9.0±0.3 | 21.1±4.7 | 2.4±0.6 | 5.3±1.8 | 6.1±5.9 | 1.7±1-4 |

<sup>1</sup>ED<sub>50</sub>, effective dose 50, the dose of compound that reduces by 50% the fluorescence (FRA) or virus titers (VIA). Data shown are mean and standard deviation of the mean relative to FRA (*left panel*) and VIA (*right panels*) described in Figure 8. The numbers between brackets indicate the number of independent experiments for each assay. <sup>2</sup>Ratio between values obtained for AD169-GFP26 and AD169-GFP.

### Identification of compounds interfering with HCMV replication

SMs were subsequently screened for their ability to interfere with HCMV replication. Each SMs was tested at two different concentrations (10 and 100 µM), and GCV (50 µM) was included as a positive control for inhibition of viral replication. After infection and compound treatment, cells were monitored by light microscopy daily for 7 days p.i. to evaluate CPE, presence of precipitates, and viral replication. At 7 days p.i., cells were lysed and the plates processed for fluorimetric quantification of the levels of viral replication relative to each condition. In parallel, MTT assays were performed with uninfected cells. At the lowest concentration tested (10 μM), only one compound (A4) was not soluble and caused evident cytotoxicity, whereas two compounds (B3 and B6) significantly inhibited viral replication (Figure 5A). On the other hand, at the highest concentration tested (100 µM) several of the 18 SMs formed visible precipitates and caused cell death, indicating poor solubility and toxicity, while two compounds (B1 and C6) impaired viral replication without markedly affecting cell viability (Figure 5B). In summary, our data show that four SMs reduced viral replication in the absence of precipitates and evident cell cytotoxicity, to levels similar to those observed upon GCV treatment. Such inhibition was confirmed by microscopic analysis of infected cells, with a noticeable decrease in CPE and number and intensity of YFP-positive cells, to similar levels as GCV-treated cells (not shown).

**Figure 5.**
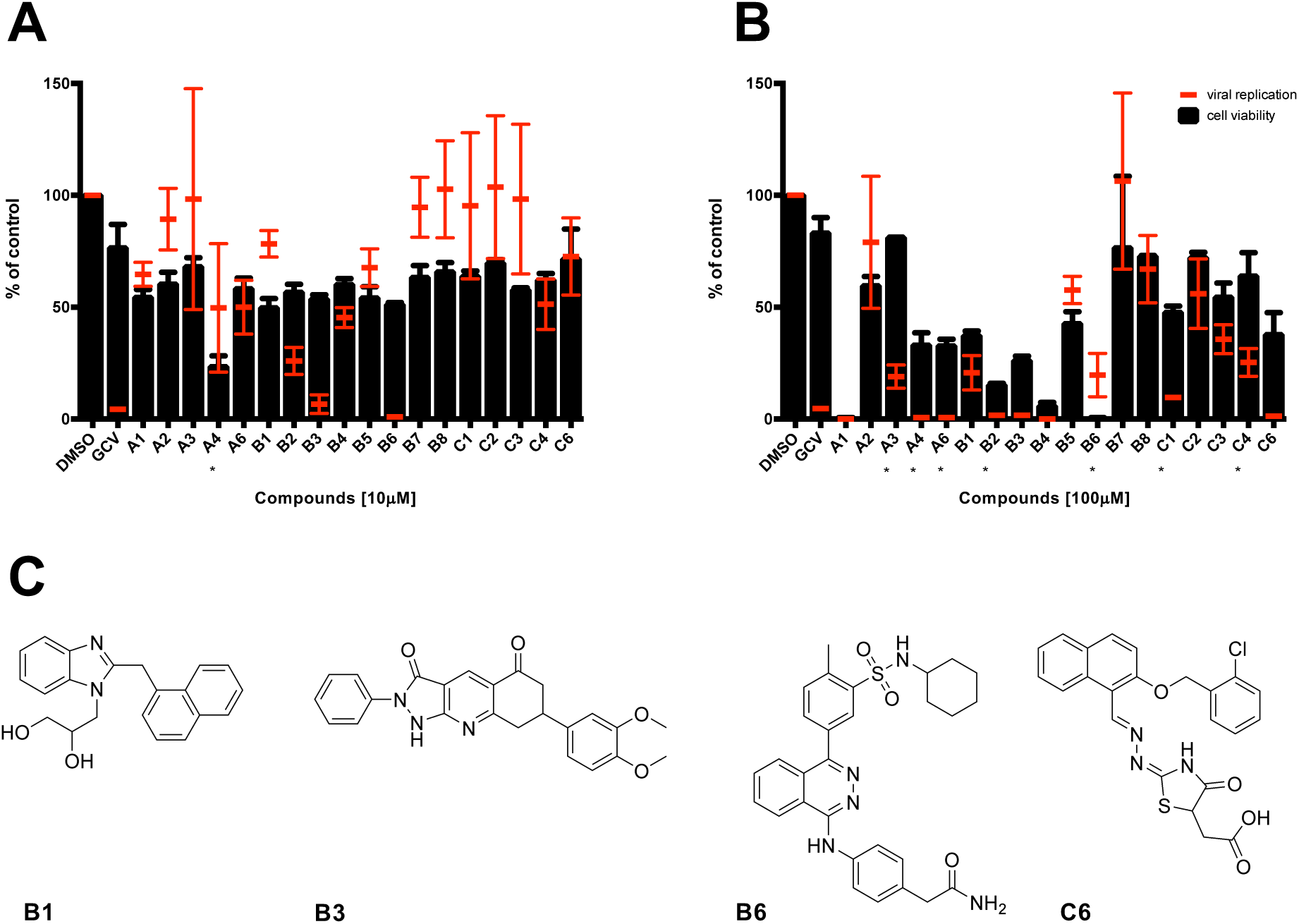
Identification of compounds interfering with HCMV replication. To assess the effect of the SMs on HCMV replication, MRC5 cells were infected with TB4-UL83-EYFP and treated with each SM either at concentration of 10 μM (A) or 100 μM (B). In parallel, uninfected MRC5 cells were also treated for assessment of SMs cytotoxicity. Seven days post treatment, cells were processed for data acquisition and analysis as described in the Materials and Methods section. Mean values YFP values relative to infected cells treated with the indicated compounds are expressed as a percentage of DMSO-treated cells (*black bars*). Cell viability was assessed by MTT assays, and data expressed as a percentage of DMSO treated cells (*red bars*). The mean + S.E.M of 3 independent experiments is shown. * = presence of precipitates. (C) The chemical structure of active molecules is shown.

### Dose-dependent inhibition of HCMV replication

We then decided to determine the ED_50_ and CC_50_ values of each of the 4 SMs identified in our small-scale FRA-based screening. To this end, we performed dose-response FRAs and MTT assays in MRC5 cells treated with increasing concentrations of each SM. Importantly, all tested SMs reproducibly inhibited viral replication in a dose-dependent manner, with ED_50_ values in the low micromolar range (Figure 6). As expected, GCV efficiently inhibited HCMV replication with a ED_50_ of 2.3±0.7 μM and did not cause detectable cytotoxic effects at any concentration tested (Figure 6). Among the SMs tested B6 was the most active compound and exhibited an ED_50_ slightly lower that that calculated for GCV (2.5±0.7 μM). However, it was also endowed with considerable cytotoxicity (CC_50_ of ∼10 μM), resulting in an SI < 5. On the other hand, B3, the second most active SM (ED_50_ of 4.2±2.4 μM), did not cause evident cytotoxicity up to 100 μM, and therefore had an SI > 20. C6 also exhibited low cytotoxicity, but was significantly less efficient in inhibiting HCMV replication than the other two SMs (ED_50_ of 16.5±10.3 μM), and cell morphology appeared significantly altered upon microscopic evaluation (not shown). Finally, the effect of B1 on HCMV life cycle was evident only at high concentrations (ED_50_ of 86.7±10.3 μM).

**Figure 6.**
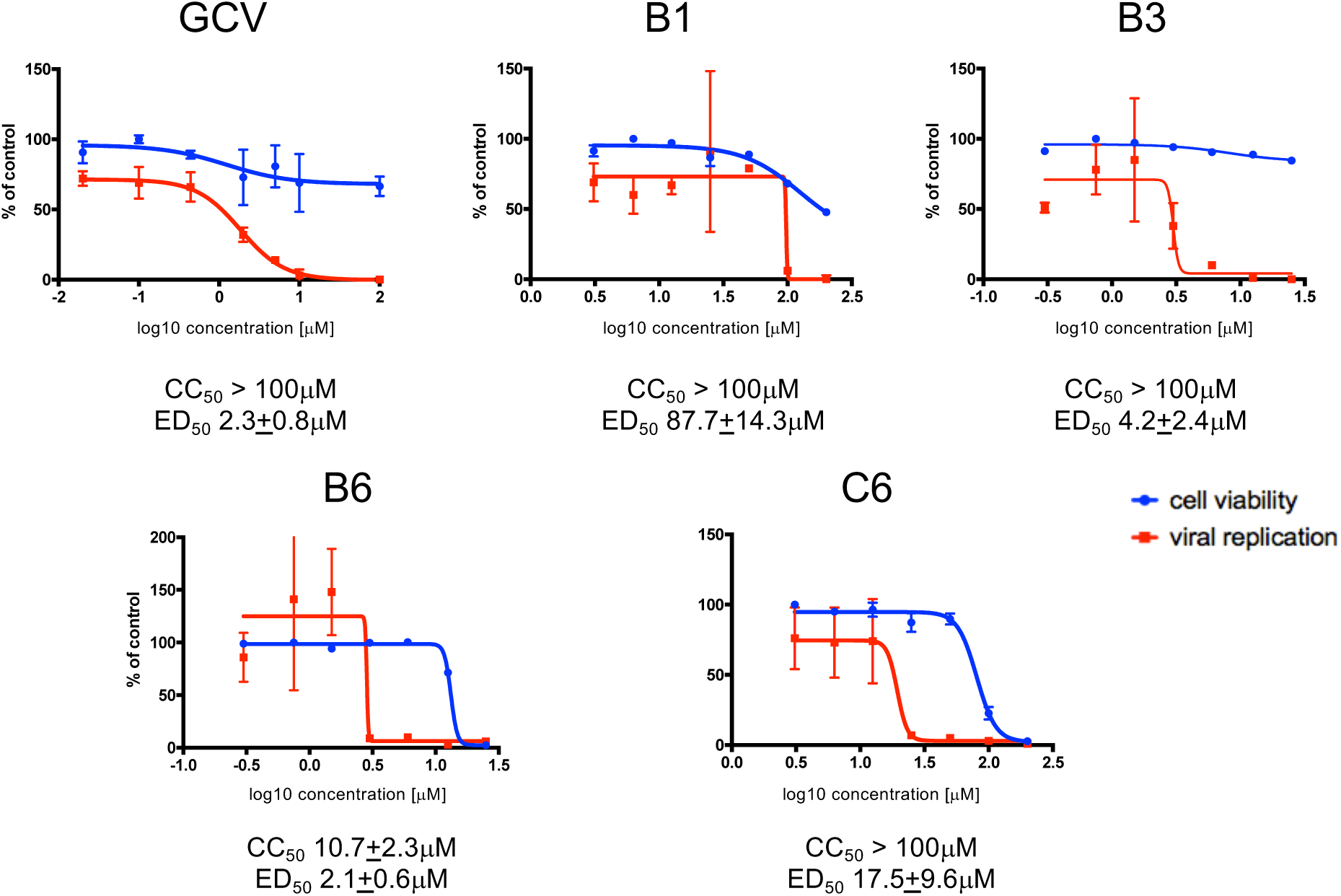
Determination of ED_50_ and CC_50_ values of SMs by FRA and MTT assay. MRC5 cells were infected with TB4-UL83-EYFP and treated with increasing concentrations of the indicated compounds. In parallel, uninfected MRC5 cells were also treated for assessment of SMs cytotoxicity. Seven days post treatment, cells were processed for data acquisition and analysis as described in the Materials and Methods section. (A) In parallel, uninfected MRC5 cells were also treated for assessment of SMs cytotoxicity. Seven days post treatment, cells were processed for data acquisition and analysis as described in the Materials and Methods section. Mean values YFP values relative to infected cells treated with the indicated compounds are expressed as a percentage of DMSO-treated cells (*red squares*). Cell viability was assessed by MTT assays, and data expressed as a percentage of DMSO treated cells (*blue circles*). For each compound, representative plots are shown, along with the cytotoxic concentration 50 (CC_50_) and effective dose 50 (ED_50_) mean values ± standard error of the mean of at least 4 independent experiments.

Overall, our data indicate that SM B3 might be specifically interfering with HCMV life cycle. Analysis of its predicted binding mode to UL44 revealed several hydrophobic interactions with UL44 residues M116, C117, A118, P119, F121, M123, L99, L93 as well as two H-bond interactions with S96 (Figure 7). The ability to B3 to inhibit HCMV AD169 replication in HFF was also assessed by PRA assays (not shown), resulting in a ED_50_ of 16.3±8.5 μM (n=4).

**Figure 7.**
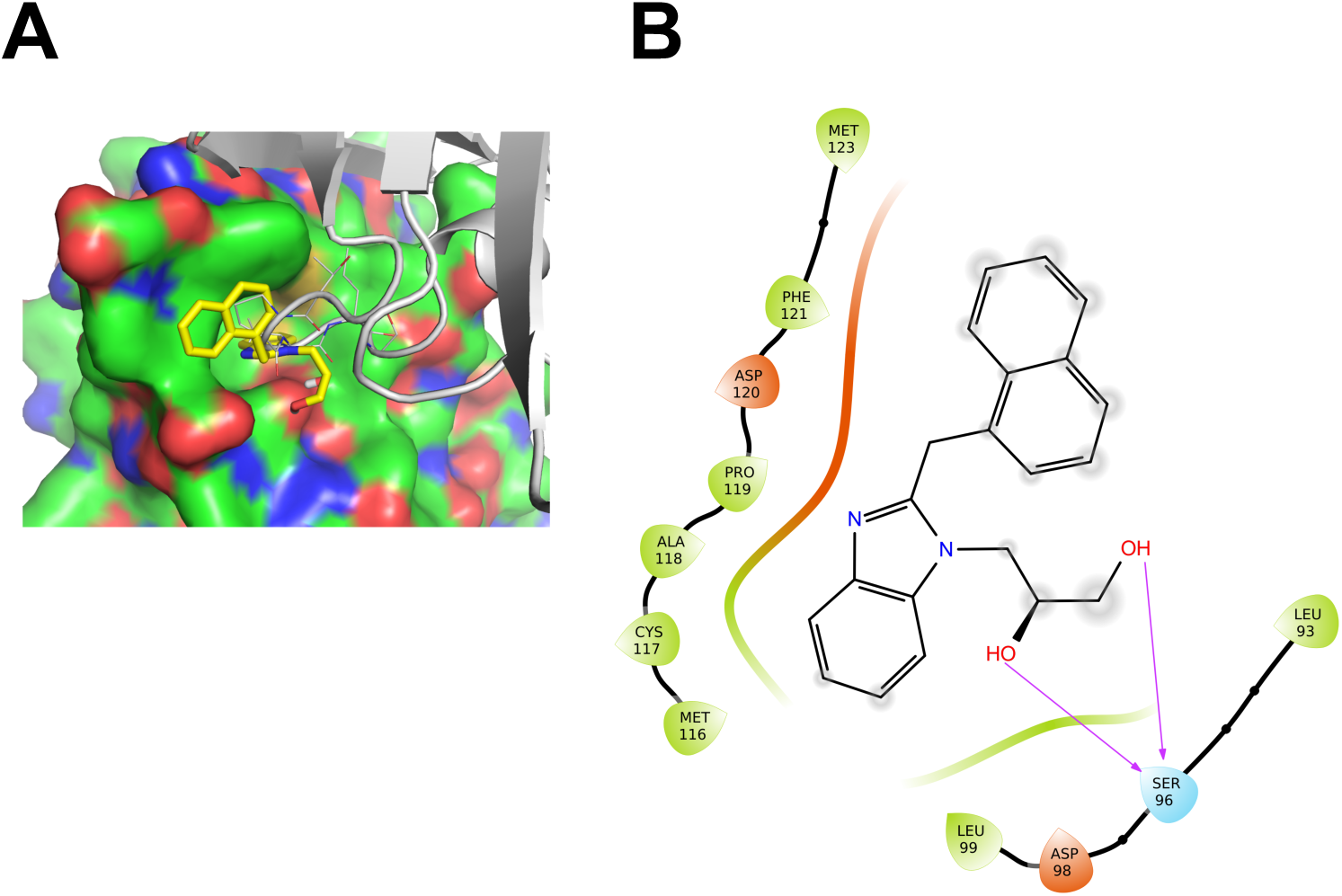
Analysis of the theoretical binding mode of B3 to UL44. A graphic representation of UL44(1-290) homodimers in the presence of B3 is shown. (A) One monomer is represented as surface, the other one as ribbons, with residues involved in the dimerization being shown as sticks, whereas B3 is shown as yellow sticks. (B) 2D ligand interaction diagram of the theoretical binding mode for B3 is shown. Hydrogen bond interactions are shown as violet arrows. Positive and negative charged amino acids are represented in blue and red, respectively. Residues involved in hydrophobic or polar interactions are shown in green and light blue, respectively. Ligand-exposed fractions are indicated as a gray, circular shadow.

### SM B3 inhibits replication of a HCMV GCV-resistant strain

An important characteristic of an antiviral interfering with UL44 dimerization would be its ability to inhibit replication of viral strains resistant to the currently used antivirals. To verify if compound B3 is endowed with such ability, we compared its ED_50_ against a recombinant reporter virus AD169-GFP and its GCV-resistant counterpart AD169-GFP26, bearing the UL97 M460I substitution (57). To this end, MRC5 cells were infected with of either AD169-GFP or AD169-GFP26 virus at MOI of 0.05 infectious units/cell, treated with B3 or GCV for 1 week, and then viral replication was assessed by FRA whereas viral titers in cell culture supernatants were simultaneously quantified by VYAs. FRA assays (Figure 7A, C and Table II) revealed that GCV inhibited more efficiently replication of the AD169-GFP virus than of AD169-GFP26 (ED_50_ of 2.6±0.8 μM and 20.0±4.3 μM, respectively; n=5), while AD169-GFP and AD169-GFP26 appeared equally sensitive to B3 (ED_50_ of 9.0±0.3 μM and 21.1±4.7 μM, respectively; n=5). Very similar results were obtained after quantification of viral progeny in VYAs (Figure 8C, D). Indeed, GCV inhibited more efficiently viral production of HCMV AD169 (ED_50_ of 0.5±0.47 μM; n=3) as compared to AD169-GFP26 (ED_50_ of 4.5±4.5 μM; n=3), whereas B3 did not (ED_50_ of 5.3+1.8 μM and 6.1+5.9 μM, respectively). Thus, B3 appears to efficiently impair replication of both GCV-sensitive and GCV-resistant HCMV.

**Figure 8.**
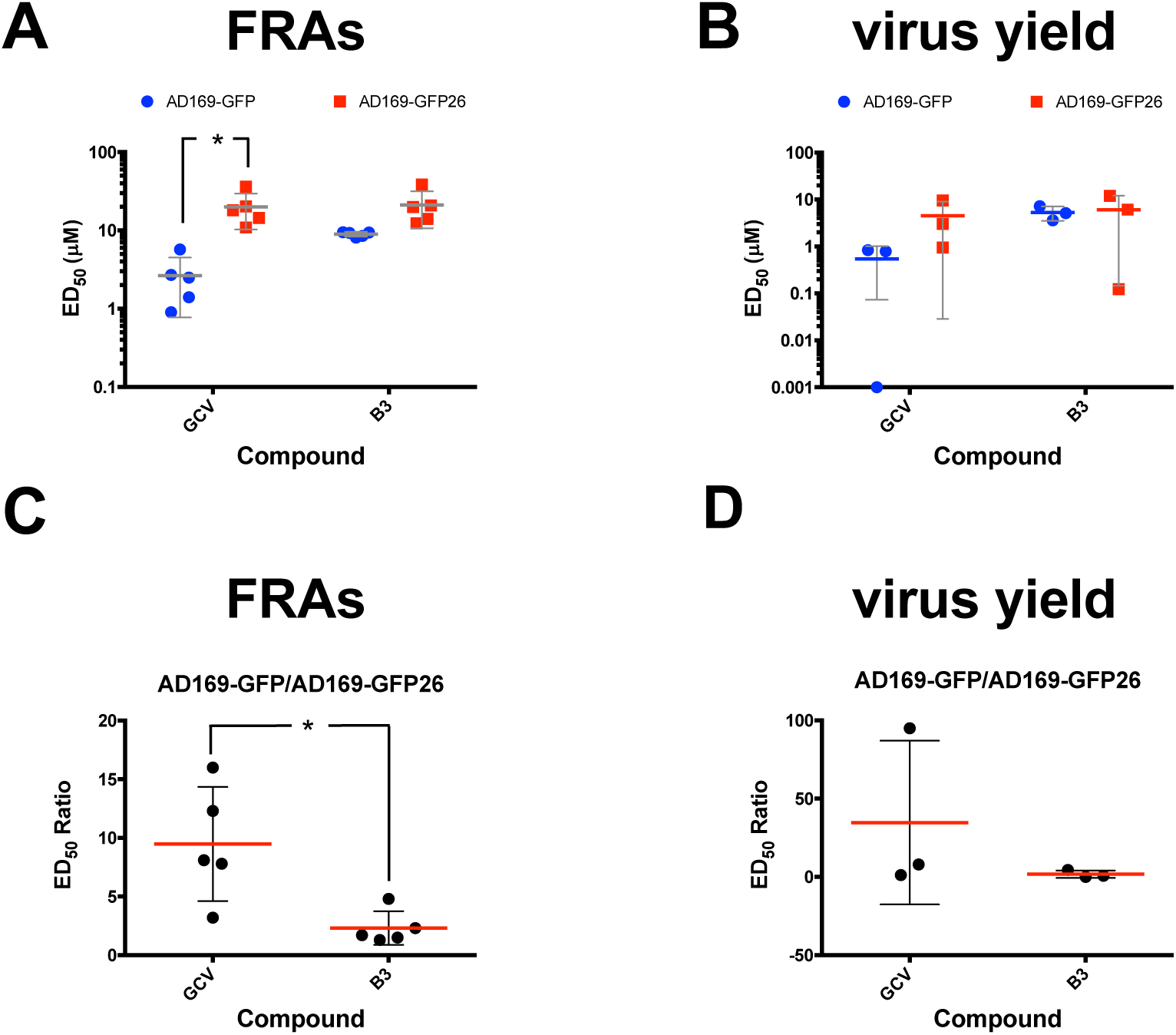
B3 efficiently inhibits replication of the GCV-resistant AD169-GFP26 virus. MRC5 were infected with either AD169-GFP virus or its GCV-resistant counterpart AD169-GFP26 treated with increasing concentrations of the indicated compounds. At 7 days p.i., cells were lysed and plates processed for fluorimetric analysis using a plate reader and supernatants collected for virus yield reduction assays (A, C). MRC5 cells were infected with serial dilutions of supernatants derived from infected cells. At 7 days p.i., viral titers were calculated using the TCID_50_ method (B, D). Data from both assays were used to calculate the ED_50_s relative to the two viruses (A, B) as well as the ratio between the ED_50_ calculated for AD169-GFP26 and AD169-GFP (C, D) relative to GCV and B3. Data shown are single measurements, means, and standard deviations of at least three independent experiments, along with the p-value relative to the Student’s t-test the indicated groups; *: p ≤ 0.05.

### B3 selectively impairs HCMV late gene expression

In order to characterize the mode of action of B3, MRC5 cells were infected with AD169 at an MOI of 1 infectious units/cell for 2 h and expression of immediate-early (IE1/2), early-late (UL44 and pp65), and late (pp28) proteins was assessed at different time points p.i. As expected, neither GCV nor B3 treatment affected IE1/2 expression at 6 and 12 h p.i., indicating that the observed decrease in viral replication did not depend on an inhibition of activity on the major IE promoter (Figure 9A, B). Similarly, UL44 and pp65 levels were not affected up to 24 h p.i. (Figure 9C), confirming that IE function was not compromised. At 48 h p.i., a decrease in the expression of UL44 and –to a greater extent-of pp65 could be observed (Figure 9D). Importantly, both GCV and B3 treatment strongly inhibited pp28 expression at both 72 h and 96 h p.i. (Figure 8E, F). Densitometric analysis confirmed that GCV (Figure 9G) and B3 (Figure 9H) inhibited HCMV gene expression with very similar kinetics, suggesting inhibition of viral DNA synthesis and interference with an early HCMV function.

**Figure 9.**
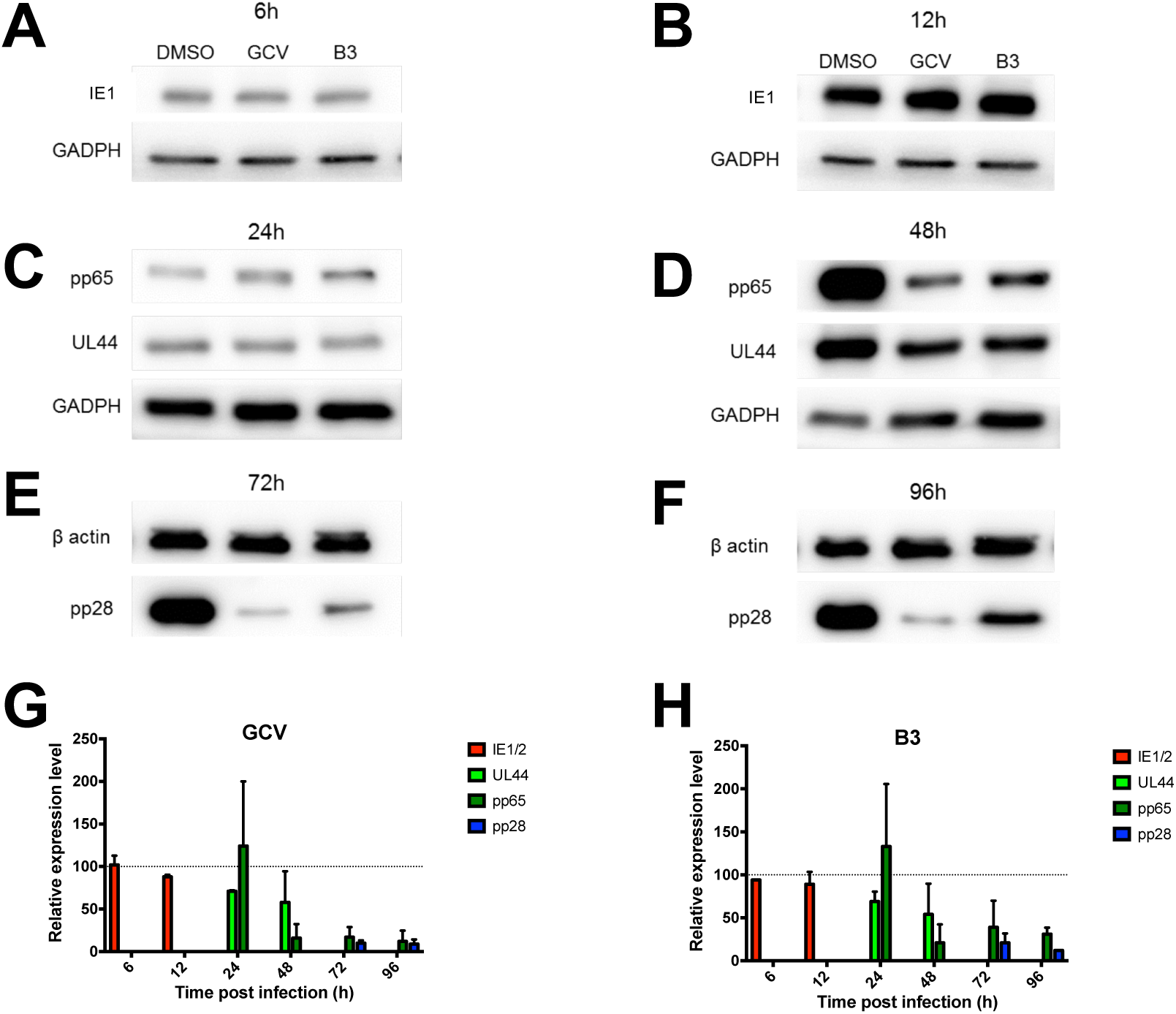
B3 specifically impairs early and late HCMV AD169 gene expression. MRC5 were infected with HCMV AD169 and treated as described in the Materials and Methods section. At the indicated time points p.i., cells were lysed and processed for Western Blotting to detect the expression of the immediate early IE1 antigen (A, B; at 6 and 12 h p.i.), the early-late antigens ppUL44 and pp65 (C, D; at 24 and 48 h p.i.), and the late antigen pp28 (E, F; at 72 and 96 h p.i.). GAPDH or β-actin were also detected as loading controls. (G, H) Loading controls were used to normalize signal intensity relative to each antigen after treatment with GCV (G) or B3 (H). Data shown are the mean ± standard deviation of the mean relative to three independent experiments.

## DISCUSSION

This is the first study directly targeting the dimerization of HCMV DNA polymerase accessory subunit UL44. The latter is an interesting therapeutic target considering the interaction interface shown in the crystal structure of UL44(1-290), and the fact that single amino acid substitutions affecting dimerization *in vitro* also impaired dsDNA binding (16) and prevented *ori*Lyt dependent DNA replication in trans-complementation assays (23). Our new data presented here further strengthened this hypothesis, by showing that the same substitutions affect dimerization of full length UL44 both in *vitro* and in living cells (see Figs. 1-4). Indeed, the L86A/L87A substitution reduced binding to GST-UL44 in pulldown assays (Figure 1A), and greatly reduced protein stability in TSA (Figure 1B). The latter observation is consistent with the fact that shown that substitutions affecting protein multimerization decrease the Tm (41). Similarly, the L86A/L87A substitution reduced FRET efficiency and the B_50_ value for UL44 self-interaction, but had little or no effect on the ability of the protein to bind to its catalytic subunit UL54 (Figures 2 and 3), therefore suggesting that the protein was still properly folded. Intriguingly, substitutions preventing nuclear translocation or DNA binding of UL44 did not prevent its dimerization or binding to the DNA polymerase catalytic subunit UL54, as exemplified by B_50_ values very similar to the wild-type protein, and consistently with our previously published model whereby UL44 dimerizes and associates with UL54 in the cytosol before nuclear import (3, 26). However, both substitutions significantly reduced the Bmax values as compared to the wild-type protein, which is possibly due to a change in protein conformation (and therefore energy transfer between BRET donor and acceptor subunits) of UL44 homodimers upon DNA binding. It is worth mentioning that, while the L86A/L87A substitutions similarly caused a decrease in the Bmax value relative to UL44 dimerization, the F121A substitution did not. This is interesting and consistent with the evidence that the F121A variant binds to DNA with a 10-fold greater affinity than the L86A/L87A variant and the hypothesis that it still presents weak DNA binding upon dimerization (16). Regardless of the effects of DNA binding on the HCMV holoenzyme conformation, which are currently being investigated in our laboratory, our data show that UL44 interactions with UL54 and with itself occur with similar affinity, as implied by the very similar B_50_ values (Figure 3 and Table I; (56)). Therefore, given the fact that previous studies have identified peptides and SMs able to successfully block HCMV replication by interfering with the interaction between UL54 and UL44 (50, 51, 65), these results prompted us to perform a virtual screening aimed at identifying SMs inhibiting UL44 dimerization (Figure 4). One of the 18 SMs tested was capable of inhibiting HCMV replication in a variety of assays, although with different efficacy. ED_50_ values ranged from 4.2 (FRA) to 16.3 μM (PRA), while those determined for GCV were comprised between 0.5 (VYA) and 2.6 (FRA) μM (Table III). The ED_50_ values calculated here for GCV are compatible with those reported previously in the literature, with some variance being attributable to intrinsic differences between the different assays and viruses tested. For example, FRA with the TB40-UL83-EYFP virus rely on the measurement of pp65 expression, which is expressed with an early-late kinetic, whereas the expression of the reporter gene in the AD169-GFP virus is under control of the IE promoter (57, 58). Importantly, B3 inhibited the AD169-GFP virus and its GCV-resistant derivative AD169-GFP26 with almost identical ED_50_ values in both FRA and virus yield reduction assays while GCV, as expected, was much more active towards AD169-GFP (see Figure 8 and Table II). Although we did not formally prove here that B3 acts by disrupting the UL44 homodimer during viral infection, the fact that it inhibits HCMV gene expression starting from 48 p.i., and strongly inhibits late gene expression at 72 and 96 h p.i., in a very similar fashion to GCV (Figure 9) is compatible with inhibition of viral DNA replication (66–68). Therefore, our results raise hopes in terms of potential use of HCMV dimerization inhibitors for the treatment of patients infected with drug-resistant HCMVs.

**Table III.**
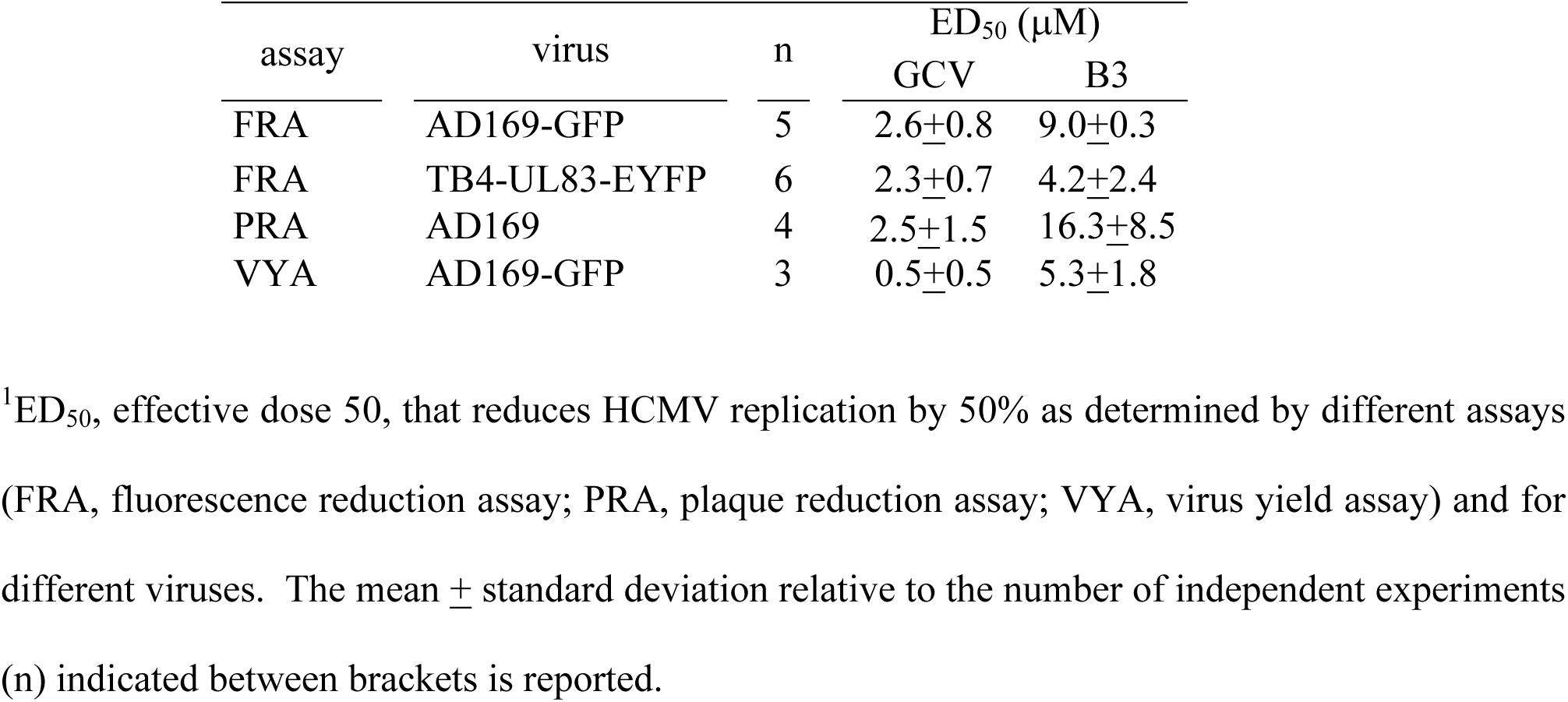
Summary of ED_50_ values calculated for GCV and B3 in this study.

| assay | virus | n | ED<sub>50</sub> (μM) |  |
| --- | --- | --- | --- | --- |
|  |  |  | GCV | B3 |
| FRA | AD169-GFP | 5 | 2.6±0.8 | 9.0±0.3 |
| FRA | TB4-UL83-EYFP | 6 | 2.3±0.7 | 4.2±2.4 |
| PRA | AD169 | 4 | 2.5±1.5 | 16.3±8.5 |
| VYA | AD169-GFP | 3 | 0.5±0.5 | 5.3±1.8 |

<sup>1</sup>ED<sub>50</sub>, effective dose 50, that reduces HCMV replication by 50% as determined by different assays
(FRA, fluorescence reduction assay; PRA, plaque reduction assay; VYA, virus yield assay) and for
different viruses. The mean ± standard deviation relative to the number of independent experiments
(n) indicated between brackets is reported.

## Acknowledgments

This study was supported by Institutional research grants (ex 60%), PRAT2013 grant CPDA130224/13 and PRID2019 grant from University of Padua to GA, by Associazione Italiana per la Ricerca sul Cancro, Italy (grant IG18855 to A.L.), by British Society for Antimicrobial Chemotherapy, UK (grant bsac-2018-0064 to A.L.), by Ministero dell’Istruzione dell’Università e della Ricerca, Italy (grant 2017KM79NN to A.L.), and by STARS Consolidator Grant FINDER (to B.M.). Recombinant TB40-UL83-EYFP virus as well as AD169-GFP and AD169-GFP26 viruses were kindly provided by Micheal Winkler (Göttingen, Germany) and Manfred Marschall (Erlangen, Germany), respectively.

## Author contributions

Designed the study: GA and FF. Performed experiments: MT, VDA, FF, HG, BM and GA. Performed the virtual screening: FF; Analyzed experiments: GA, BM, MT, VDA and HG. Supervised the study: GA, AL, and GP. Wrote the manuscript: HG, FF and GA. Revised the manuscript: GA, AL, BM and HG.

## APPENDIX

### Viruses, viral stocks preparation and titration

For viral stocks preparation, MRC5 cells were seeded in T150 flasks (6×10^6^ cells/flask) the day before infection, which was carried in the absence of serum, at a multiplicity of infection (MOI) of 0.02-0.05 infectious units/cell, for 2 h in a humidified incubator at 37°C and 5% CO_2_ with occasional shaking. Cell medium was changed at the end of the incubation time and every 3 days. Viral supernatants were collected 7 days p.i., centrifuged for 5’ at 700 rpm to remove cell debris, and stored in the -80°C after addition of 1% (v/v) DMSO. For titration of infectivity by immunological detection of IE proteins, MRC5 cells were seeded in 96 well plates (1.5×10^4^ cells/well) in 200 μl of DMEM/well. The day after, viral stocks were rapidly thawed at 37°C as described above. Serial 10-fold dilutions of viral stocks were used to infect three wells per dilution for 2 h at 37°C in a final volume of 200 μl. Twenty for h later, media was removed and cells washed once with PBS (200 μl/well). Subsequently cells were fixed for 15’ with EtOh 95% at RT followed by saturation of unspecific binding sites by incubation with PBS containing 5% FBS (w/v) for 1h at 37°C (50 μl/well). After 3 washes with PBS, cells were incubated with *α*-IE1&2 mAb (Virusys Corporation, #CA003-1; 1:100) diluted in PBS containing 5% FBS for 16 h at +4°C. Cells were washed 3x with PBS and incubated with either Alexa Fluor 488 Alexa Fluor 555 the goat anti-mouse IgG secondary antibodies (Themofisher, #A-11001 and #A-21424, respectively). After 3x washes with PBS positive cells were visualized and counted using an inverted fluorescent microscope (Leica, DFC420 C), to allow calculation of the viral titer expressed as infectious units/ml.

### Proteins Expression and Purification

Cells were grown in LB medium containing 100 µg/ml ampicillin until the OD_600_ was 0.8 and then induced by the addition of 0.3 mM IPTG for 3 h at room temperature (RT). Cells were pelleted, resuspended in lysis buffer (50 mM NaH_2_PO_4_, 500 mM NaCl, 10% glycerol, 10 mM imidazole, 1 mg/ml lysozyme, and Complete protease inhibitors), and then lysed by two freeze/thaw cycles followed by sonication. The lysate was centrifuged at 13,000 rpm for 45 min, applied to a 0.5-ml Ni-NTA agarose resin column (Qiagen) that had been equilibrated in lysis buffer, and then washed with wash buffer 1 (50 mM NaH_2_PO_4_, 500 mM NaCl, 10% glycerol, and 20 mM imidazole) and subsequently with wash buffer 2 (50 mM NaH_2_PO_4_, 500 mM NaCl, 10% glycerol, and 50 mM imidazole). Finally, proteins were eluted with elution buffer (50 mM NaH_2_PO_4_, 500 mM NaCl, 10% glycerol, and 250 mM imidazole). Purified proteins were dialyzed against 20 mM Tris-HCl pH 7.5, 150 mM NaCl, 30% glycerol, 0.1 mM EDTA, 2 mM DTT and stored at -80°C.

### Western-blot assays

Samples were diluted in Laemmli sample buffer (0.05 M Tris-HCl, pH 6.8, 0.05% Bromophenol blue, 0.1 M DTT, 10% Glycerol [v/v], 2% SDS) and boiled 5 minutes at 95°C before being loaded and electrophoretically separated on 8.5 % polyacrylamide gels. Separated proteins were blotted on polyvinylidene fluoride (PVDF) membranes (GE Healthcare, #RPN303F). Membranes were saturated with PBS containing 0.2% Tween20 and 5% milk (w/v), and incubated with the appropriate primary and secondary antibodies diluted in PBS containing 0.2% Tween20 and 5% milk (w/v). For analysis of fractions deriving from GST-pulldown assays, membranes were incubated with an enhanced chemiluminescence substrate (LiteAblot Extend Long Lasting Chemiluminescent Substrate Kit, Euroclone) and signal acquired using an imaging system (Versadoc, Biorad). For analysis of cell lysates, membranes were incubated with an enhanced chemiluminescence substrate (ECL Prime Western Blotting Detection Reagent, GE Healthcare, #RPN2236).

### Cell cytotoxicity assays

At the desired time point post treatment, 10 µl of MTT (5 mg/ml in PBS) were added to each well of the 96 well plate, and cells were further incubated at 37°C with 5% CO_2_ for 2 h. Subsequently, cells were lysed by gently pipetting in 100 µl of Lysis Buffer (SDS 10%, 0.01M HCl in H_2_0). Plates were then incubated overnight at 37°C. Absorbance was determined with a spectrophotometer plate reader at 620 nm (Sunrise, Tecan). Data were exported to excel and background signal relative to the average of DMEM only containing wells were subtracted from each well. For calculation of 50% cytotoxic concentration (CC_50_), defined as the concentration of compound required to decrease cell metabolic activity by 50%, data were exported to Graphpad Prism (Graphpad Software Inc.), *x* data transformed to logarithmic and fitted to a linear regression or to a 4 Parameter Logisitic Equation (Sigmoidal).

### Bioluminescence resonance energy transfer (BRET) assays

HEK 293T cells were seeded in a 24-well plate (1×10^5^ cells/well) and the next day were transfected using Lipofectamin 2000 (Life Technologies) following the manufacturer’s recommendations, with appropriate amounts of BRET donor and BRET acceptor expressing plasmids. For each construct, the donor (RLuc) expressing plasmid was transfected both in the absence and in the presence of the relative acceptor (YFP) expressing plasmid to allow calculation of background BRET signal. At 48 h post-transfection, culture medium was removed from wells and cells were gently washed with 1 ml of PBS, before being resuspended in 290 µl of PBS. Cell resuspensions (90 µl) were then transferred to a 96-well black flat bottom polystyrene TC-treated microplate (Costar, #3916) in triplicate and signals were acquired using a reader compatible with BRET measurements (VICTOR X2 Multilabel Plate Reader, Perkin Elmer). Fluorescent signals (YFPnet) relative to YFP fluorescent emission were acquired using a fluorimetric excitation filter (band pass 485 ± 14 nm) and a fluorimetric emission filter (band pass 535 ± 25 nm). Luminometric readings were performed at 5’, 15’, 30’, 45’ and 60’ after addition of native Coelenterazine (5 μM final concentration, PJK). Data were acquired for 1 sec/well, using a luminometric 535 ± 25 nm emission filter (YFP signal) and a luminometric 460 ± 25 nm emission filter (RLuc signal). Before reading, the plate was shaken for 1 sec. at normal speed and with double orbit. After background subtraction using values relative to mock transfected cells, the data obtained were used to calculate the BRET signal, defined as the ratio between the YFP and RLuc signals calculated for a specific BRET pair, according to the formula:

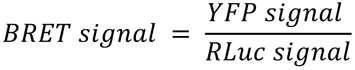

Similarly, the BRET ratio, defined as the difference between the BRET value relative to a BRET pair and the BRET value relative to the BRET donor alone, was calculated according to the formula:

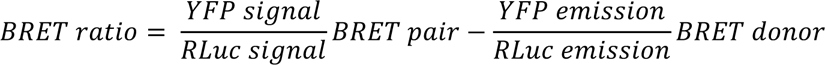

### Fluorescence resonant energy transfer (FRET) acceptor photobleaching assays

HEK 293T cells were seeded in a 24-well plate onto glass coverslips (1×10^5^ cells/well) and the next day were transfected with 125 and 250 ng of YFP and CFP expression constructs, respectively, using Lipofectamine 2000 (Life Technologies) following the manufacturer’s recommendations. At 48 h post-transfection, cells were washed with PBS and fixed with 4% paraformaldehyde for 10’ at room temperature. After a wash with milliQ water, the samples were mounted on glass slides with Fluoromount G (Southern Biotech). Samples were imaged by CLSM using a Leica SP2 confocal laser scanning microscope (Leica) equipped with a 63× oil immersion objective. FRET efficiency was determined according to the following formula: FRETeff = [(EDpost − EDpre)/EDpost] × 100, where ED represents the emitted donor fluorescence before (EDpre) or after (EDpost) photobleaching of the acceptor fluorophore. Measurements of at least 20 cells from two independent transfections were used to calculate the box plots. Significance values were calculated by using the unpaired t test with the GraphPad Prism 6 software package (Graphpad Software Inc.). Images were processed using ImageJ (NIH).

## REFERENCES

1. Griffiths P, Baraniak I, Reeves M. 2015. The pathogenesis of human cytomegalovirus. J Pathol 235:288–297.

2. Britt WJ. 2018. Maternal Immunity and the Natural History of Congenital Human Cytomegalovirus Infection. Viruses 10.

3. Alvisi G, Jans DA, Camozzi D, Avanzi S, Loregian A, Ripalti A, Palu G. 2013. Regulated transport into the nucleus of herpesviridae DNA replication core proteins. Viruses 5:2210–2234.

4. Ertl PF, Powell KL. 1992. Physical and functional interaction of human cytomegalovirus DNA polymerase and its accessory protein (ICP36) expressed in insect cells. J Virol 66:4126–4133.

5. De Clercq E. 2013. Selective anti-herpesvirus agents. Antivir Chem Chemother 23:93–101.

6. Lurain NS, Chou S. 2010. Antiviral drug resistance of human cytomegalovirus. Clin Microbiol Rev 23:689–712.

7. Drouot E, Piret J, Lebel MH, Boivin G. 2014. Characterization of multiple cytomegalovirus drug resistance mutations detected in a hematopoietic stem cell transplant recipient by recombinant phenotyping. J Clin Microbiol 52:4043–4046.

8. Goldner T, Hewlett G, Ettischer N, Ruebsamen-Schaeff H, Zimmermann H, Lischka P. 2011. The novel anticytomegalovirus compound AIC246 (Letermovir) inhibits human cytomegalovirus replication through a specific antiviral mechanism that involves the viral terminase. J Virol 85:10884–10893.

9. Kim ES. 2018. Letermovir: First Global Approval. Drugs 78:147–152.

10. Jung S, Michel M, Stamminger T, Michel D. 2019. Fast breakthrough of resistant cytomegalovirus during secondary letermovir prophylaxis in a hematopoietic stem cell transplant recipient. BMC Infect Dis 19:388.

11. Lischka P, Michel D, Zimmermann H. 2016. Characterization of Cytomegalovirus Breakthrough Events in a Phase 2 Prophylaxis Trial of Letermovir (AIC246, MK 8228). J Infect Dis 213:23–30.

12. Ripalti A, Boccuni MC, Campanini F, Landini MP. 1995. Cytomegalovirus-mediated induction of antisense mRNA expression to UL44 inhibits virus replication in an astrocytoma cell line: identification of an essential gene. J Virol 69:2047–2057.

13. Gallo ML, Dorsky DI, Crumpacker CS, Parris DS. 1989. The essential 65-kilodalton DNA-binding protein of herpes simplex virus stimulates the virus-encoded DNA polymerase. J Virol 63:5023–5029.

14. Gibson W, Murphy TL, Roby C. 1981. Cytomegalovirus-infected cells contain a DNA-binding protein. Virology 111:251–262.

15. Weiland KL, Oien NL, Homa F, Wathen MW. 1994. Functional analysis of human cytomegalovirus polymerase accessory protein. Virus Res 34:191–206.

16. Appleton BA, Loregian A, Filman DJ, Coen DM, Hogle JM. 2004. The cytomegalovirus DNA polymerase subunit UL44 forms a C clamp-shaped dimer. Mol Cell 15:233–244.

17. Boccuni MC, Campanini F, Battista MC, Bergamini G, Dal Monte P, Ripalti A, Landini MP. 1998. Human cytomegalovirus product UL44 downregulates the transactivation of HIV-1 long terminal repeat. AIDS 12:365–372.

18. Silva LA, Loregian A, Pari GS, Strang BL, Coen DM. 2010. The carboxy-terminal segment of the human cytomegalovirus DNA polymerase accessory subunit UL44 is crucial for viral replication. J Virol 84:11563–11568.

19. Silva LA, Strang BL, Lin EW, Kamil JP, Coen DM. 2011. Sites and roles of phosphorylation of the human cytomegalovirus DNA polymerase subunit UL44. Virology 417:268–280.

20. Alvisi G, Jans D, Guo J, Pinna L, Ripalti A. 2005. A protein kinase CK2 site flanking the nuclear targeting signal enhances nuclear transport of human cytomegalovirus ppUL44. Traffic 6:1002–1013.

21. Alvisi G, Marin O, Pari G, Mancini M, Avanzi S, Loregian A, Jans DA, Ripalti A. 2011. Multiple phosphorylation sites at the C-terminus regulate nuclear import of HCMV DNA polymerase processivity factor ppUL44. Virology 417:259–267.

22. Loregian A, Appleton BA, Hogle JM, Coen DM. 2004. Specific residues in the connector loop of the human cytomegalovirus DNA polymerase accessory protein UL44 are crucial for interaction with the UL54 catalytic subunit. J Virol 78:9084–9092.

23. Sinigalia E, Alvisi G, Mercorelli B, Coen DM, Pari GS, Jans DA, Ripalti A, Palu G, Loregian A. 2008. Role of homodimerization of human cytomegalovirus DNA polymerase accessory protein UL44 in origin-dependent DNA replication in cells. J Virol 82:12574–12579.

24. Komazin-Meredith G, Petrella RJ, Santos WL, Filman DJ, Hogle JM, Verdine GL, Karplus M, Coen DM. 2008. The human cytomegalovirus UL44 C clamp wraps around DNA. Structure 16:1214–1225.

25. Jiang C, Hwang YT, Wang G, Randell JCW, Coen DM, Hwang CBC. 2007. Herpes simplex virus mutants with multiple substitutions affecting DNA binding of UL42 are impaired for viral replication and DNA synthesis. Journal of Virology 81:12077–12079.

26. Alvisi G, Jans D, Ripalti A. 2006. Human cytomegalovirus (HCMV) DNA polymerase processivity factor ppUL44 dimerizes in the cytosol before translocation to the nucleus. Biochemistry 45:6866–6872.

27. Alvisi G, Roth DM, Camozzi D, Pari GS, Loregian A, Ripalti A, Jans DA. 2009. The flexible loop of the human cytomegalovirus DNA polymerase processivity factor ppUL44 is required for efficient DNA binding and replication in cells. J Virol 83:9567–9576.

28. Zarrouk K, Piret J, Boivin G. 2017. Herpesvirus DNA polymerases: Structures, functions and inhibitors. Virus Res doi:10.1016/j.virusres.2017.01.019.

29. Arkin MR, Tang Y, Wells JA. 2014. Small-molecule inhibitors of protein-protein interactions: progressing toward the reality. Chem Biol 21:1102–1114.

30. Shimba N, Nomura AM, Marnett AB, Craik CS. 2004. Herpesvirus protease inhibition by dimer disruption. J Virol 78:6657–6665.

31. Clackson T, Wells JA. 1995. A hot spot of binding energy in a hormone-receptor interface. Science 267:383–386.

32. Loregian A, Marsden HS, Palu G. 2002. Protein-protein interactions as targets for antiviral chemotherapy. Rev Med Virol 12:239–262.

33. Celegato M, Messa L, Goracci L, Mercorelli B, Bertagnin C, Spyrakis F, Suarez I, Cousido-Siah A, Trave G, Banks L, Cruciani G, Palu G, Loregian A. 2019. A novel small-molecule inhibitor of the human papillomavirus E6-p53 interaction that reactivates p53 function and blocks cancer cells growth. Cancer Lett doi:10.1016/j.canlet.2019.10.046.

34. Muratore G, Goracci L, Mercorelli B, Foeglein A, Digard P, Cruciani G, Palu G, Loregian A. 2012. Small molecule inhibitors of influenza A and B viruses that act by disrupting subunit interactions of the viral polymerase. Proc Natl Acad Sci U S A 109:6247–6252.

35. Clackson T, Ultsch MH, Wells JA, de Vos AM. 1998. Structural and functional analysis of the 1:1 growth hormone:receptor complex reveals the molecular basis for receptor affinity. J Mol Biol 277:1111–1128.

36. Kozakov D, Hall DR, Chuang GY, Cencic R, Brenke R, Grove LE, Beglov D, Pelletier J, Whitty A, Vajda S. 2011. Structural conservation of druggable hot spots in protein-protein interfaces. Proc Natl Acad Sci U S A 108:13528–13533.

37. London N, Raveh B, Schueler-Furman O. 2013. Druggable protein-protein interactions--from hot spots to hot segments. Curr Opin Chem Biol 17:952–959.

38. Perkins JR, Diboun I, Dessailly BH, Lees JG, Orengo C. 2010. Transient protein-protein interactions: structural, functional, and network properties. Structure 18:1233–1243.

39. Basse MJ, Betzi S, Bourgeas R, Bouzidi S, Chetrit B, Hamon V, Morelli X, Roche P. 2013. 2P2Idb: a structural database dedicated to orthosteric modulation of protein-protein interactions. Nucleic Acids Res 41:D824–827.

40. Smith MC, Gestwicki JE. 2012. Features of protein-protein interactions that translate into potent inhibitors: topology, surface area and affinity. Expert Rev Mol Med 14:e16.

41. Seetoh WG, Abell C. 2016. Disrupting the Constitutive, Homodimeric Protein-Protein Interface in CK2beta Using a Biophysical Fragment-Based Approach. J Am Chem Soc 138:14303–14311.

42. Frutos S, Rodriguez-Mias RA, Madurga S, Collinet B, Reboud-Ravaux M, Ludevid D, Giralt E. 2007. Disruption of the HIV-1 protease dimer with interface peptides: structural studies using NMR spectroscopy combined with [2-(13)C]-Trp selective labeling. Biopolymers 88:164–173.

43. Bannwarth L, Reboud-Ravaux M. 2007. An alternative strategy for inhibiting multidrug-resistant mutants of the dimeric HIV-1 protease by targeting the subunit interface. Biochem Soc Trans 35:551–554.

44. Alvisi G, Ripalti A, Ngankeu A, Giannandrea M, Caraffi SG, Dias MM, Jans DA. 2006. Human cytomegalovirus DNA polymerase catalytic subunit pUL54 possesses independently acting nuclear localization and ppUL44 binding motifs. Traffic 7:1322–1332.

45. Scaturro P, Trist IM, Paul D, Kumar A, Acosta EG, Byrd CM, Jordan R, Brancale A, Bartenschlager R. 2014. Characterization of the mode-of-action of a potent Dengue virus capsid inhibitor. J Virol 88:11540–11555.

46. Smith KM, Di Antonio V, Bellucci L, Thomas DR, Caporuscio F, Ciccarese F, Ghassabian H, Wagstaff KM, Forwood JK, Jans DA, Palu G, Alvisi G. 2018. Contribution of the residue at position 4 within classical nuclear localization signals to modulating interaction with importins and nuclear targeting. Biochim Biophys Acta 1865:1114–1129.

47. Paul D, Romero-Brey I, Gouttenoire J, Stoitsova S, Krijnse-Locker J, Moradpour D, Bartenschlager R. 2011. NS4B self-interaction through conserved C-terminal elements is required for the establishment of functional hepatitis C virus replication complexes. J Virol 85:6963–6976.

48. Loregian A, Appleton BA, Hogle JM, Coen DM. 2004. Residues of human cytomegalovirus DNA polymerase catalytic subunit UL54 that are necessary and sufficient for interaction with the accessory protein UL44. J Virol 78:158–167.

49. Sinigalia E, Alvisi G, Segre CV, Mercorelli B, Muratore G, Winkler M, Hsiao HH, Urlaub H, Ripalti A, Chiocca S, Palu G, Loregian A. 2012. The human cytomegalovirus DNA polymerase processivity factor UL44 is modified by SUMO in a DNA-dependent manner. PLoS One 7:e49630.

50. Loregian A, Rigatti R, Murphy M, Schievano E, Palu G, Marsden HS. 2003. Inhibition of human cytomegalovirus DNA polymerase by C-terminal peptides from the UL54 subunit. J Virol 77:8336–8344.

51. Chen H, Coseno M, Ficarro SB, Mansueto MS, Komazin-Meredith G, Boissel S, Filman DJ, Marto JA, Hogle JM, Coen DM. 2017. A Small Covalent Allosteric Inhibitor of Human Cytomegalovirus DNA Polymerase Subunit Interactions. ACS Infect Dis 3:112–118.

52. Loregian A, Sinigalia E, Mercorelli B, Palu G, Coen DM. 2007. Binding parameters and thermodynamics of the interaction of the human cytomegalovirus DNA polymerase accessory protein, UL44, with DNA: implications for the processivity mechanism. Nucleic Acids Res 35:4779–4791.

53. Huynh K, Partch CL. 2015. Analysis of protein stability and ligand interactions by thermal shift assay. Curr Protoc Protein Sci 79:28 29 21-14.

54. King AC, Woods M, Liu W, Lu Z, Gill D, Krebs MR. 2011. High-throughput measurement, correlation analysis, and machine-learning predictions for pH and thermal stabilities of Pfizer-generated antibodies. Protein Sci 20:1546–1557.

55. Trevisan M, Di Antonio V, Radeghieri A, Palu G, Ghildyal R, Alvisi G. 2018. Molecular Requirements for Self-Interaction of the Respiratory Syncytial Virus Matrix Protein in Living Mammalian Cells. Viruses 10.

56. Sun S, Yang X, Wang Y, Shen X. 2016. In Vivo Analysis of Protein-Protein Interactions with Bioluminescence Resonance Energy Transfer (BRET): Progress and Prospects. Int J Mol Sci 17.

57. Marschall M, Freitag M, Weiler S, Sorg G, Stamminger T. 2000. Recombinant green fluorescent protein-expressing human cytomegalovirus as a tool for screening antiviral agents. Antimicrob Agents Chemother 44:1588–1597.

58. Straschewski S, Warmer M, Frascaroli G, Hohenberg H, Mertens T, Winkler M. 2010. Human cytomegaloviruses expressing yellow fluorescent fusion proteins--characterization and use in antiviral screening. PLoS One 5:e9174.

59. Britt WJ. 2010. Human cytomegalovirus: propagation, quantification, and storage. Curr Protoc Microbiol **Chapter 14:**Unit 14E 13.

60. Halgren TA. 2009. Identifying and characterizing binding sites and assessing druggability. J Chem Inf Model 49:377–389.

61. Falchi F, Caporuscio F, Recanatini M. 2014. Structure-based design of small-molecule protein-protein interaction modulators: the story so far. Future Med Chem 6:343–357.

62. Bender A, Mussa HY, Glen RC, Reiling S. 2004. Similarity searching of chemical databases using atom environment descriptors (MOLPRINT 2D): evaluation of performance. J Chem Inf Comput Sci 44:1708–1718.

63. Hierholzer JC, Killington RA. 1996. 2 - Virus isolation and quantitation, p 25–46. In Mahy BWJ, Kangro HO (ed), Virology Methods Manual doi:https://doi.org/10.1016/B978-012465330-6/50003-8. Academic Press, London.

64. Campbell RE, Tour O, Palmer AE, Steinbach PA, Baird GS, Zacharias DA, Tsien RY. 2002. A monomeric red fluorescent protein. Proc Natl Acad Sci U S A 99:7877–7882.

65. Loregian A, Coen DM. 2006. Selective anti-cytomegalovirus compounds discovered by screening for inhibitors of subunit interactions of the viral polymerase. Chem Biol 13:191–200.

66. Hahn F, Hutterer C, Henry C, Hamilton ST, Strojan H, Kraut A, Schulte U, Schutz M, Kohrt S, Wangen C, Pfizer J, Coute Y, Rawlinson WD, Strobl S, Marschall M. 2018. Novel cytomegalovirus-inhibitory compounds of the class pyrrolopyridines show a complex pattern of target binding that suggests an unusual mechanism of antiviral activity. Antiviral Res 159:84–94.

67. He R, Forman M, Mott BT, Venkatadri R, Posner GH, Arav-Boger R. 2013. Unique and highly selective anticytomegalovirus activities of artemisinin-derived dimer diphenyl phosphate stem from combination of dimer unit and a diphenyl phosphate moiety. Antimicrob Agents Chemother 57:4208–4214.

68. Weekes MP, Tomasec P, Huttlin EL, Fielding CA, Nusinow D, Stanton RJ, Wang EC, Aicheler R, Murrell I, Wilkinson GW, Lehner PJ, Gygi SP. 2014. Quantitative temporal viromics: an approach to investigate host-pathogen interaction. Cell 157:1460–1472.

